# CCmed: cross-condition mediation analysis for identifying robust trans-eQTLs and assessing their effects on human traits

**DOI:** 10.1101/803106

**Authors:** Fan Yang, Kevin J. Gleason, Jiebiao Wang, The GTEx consortium, Jubao Duan, Xin He, Brandon L Pierce, Lin S Chen

## Abstract

Trans-eQTLs collectively explain a substantial proportion of expression variation, yet are challenging to detect and replicate since their effects are individually weak. Many trans-effects are mediated by cis-gene expression and some of those effects are shared across tissue types/conditions. To detect robust cis-mediated trans-associations at the gene-level and for specific single nucleotide polymorphisms (SNPs), we proposed two Cross-Condition Mediation methods – CCmed_gene_ and CCmed_GWAS_, respectively. We analyzed data from 13 brain tissue types from the Genotype-Tissue Expression (GTEx) project, and identified trios with cis-eQTLs of a cis-gene having associations with a trans-gene, many of which show evidence of replication in other datasets. By applying CCmed_GWAS_, we identified trans-genes associated with known schizophrenia susceptibility loci. We further conducted validation analyses assessing the schizophrenia-risk-associations of the identified trans-genes, by harnessing GWAS summary statistics from the Psychiatric Genomics Consortium and multitissue eQTL statistics from GTEx.

## Introduction

The impact of genetic variation on the human transcriptome is well-established [1, 2, 3, 4]. To date, the majority of known expression quantitative trait loci (eQTLs) affect expression of local genes (cis-eQTLs) [5]. It is known that genetic variation may also affect distal or inter-chromosomal gene expression levels as trans-eQTLs. Cis-eQTLs are often defined as the SNPs within 1 Mb from the gene transcriptional start site while trans-eQTLs are defined as the SNPs beyond or on different chromosomes. It is still challenging to detect and replicate trans-eQTLs with existing eQTL data. The reason is multifaceted: though trans-eQTLs collectively explain a substantial proportion of expression variation in the genome, their effects are individually weak; other than whole-blood [6], the available sample sizes of eQTL data from specific tissue/cell types are generally limited; many identified trans-genetic effects act in a tissue-specific or cell-type-specific manner; and the significance criteria for detecting trans-associations are more stringent due to the increase in multiple testing burden.

In contrast to directly testing for trans-associations (i.e., SNPs → trans-gene) in a single or multiple tissue types, more recently an alternative and complementary strategy has been proposed to detect trans-associations mediated by cis-gene expression levels (i.e., SNPs → cis-gene → trans-gene) [7, 8, 9, 10]. It has been shown in the causal mediation literature that while directly testing for total trans-association effects is more powerful in detecting large effects of SNPs on trans-genes, mediation-based association tests are more powerful for detecting small effects when one or more mediators are measured and accounted for [11]. Studies have shown that many cis-eQTLs also affect distal gene expression [8], and a large proportion of trans-eQTL effects are at least partially mediated by cis-gene expression levels [7]. That is, trans-associations mediated by cis-gene expression are quite abundant in the genome. Moreover, by summarizing the results from single-tissue mediation analysis, it is also observed that many cis “hub” genes (with cis-eQTLs) may affect multiple trans-genes in functionally related tissue types [7], suggesting the existence and even prevalence of cross-tissue effects for cis-mediated trans-associations (i.e., trans-associations mediated by cis-gene expression).

In this work, we first propose a Cross-Condition Mediation (CCmed) method (Figure 1) to detect cis-mediated trans-associations and to boost power by jointly analyzing multiple tissue types. To establish cis-mediated trans-effects, two conditions need to be satisfied simultaneously – non-zero cis-associations and non-zero conditional correlations of cis- and trans-gene expression levels conditioning on the eQTL genotypes. Cis-association effects are often shared across many tissue types, and gene expression levels also tend to have shared correlations among related tissue types with similar cell type compositions [5]. Therefore, we argue that cis-mediated trans-association effects are also likely to be shared across related tissue types and potentially be replicated across studies. Moreover, in CCmed_gene_, we propose to study gene-level trans-associations across conditions, i.e., we consider all cis-eQTLs for a specific gene as a set and aim to detect their joint association and mediation effects on a trans-gene. Studying gene-level trans-associations would increase the chance of detecting and replicating trans-associations that are individually weak but collectively strong and robust, while also minimizing the potential impact of different genotyping platforms in different studies. We applied the CCmed_gene_ method to data from 13 brain tissue types of the GTEx project (V8) [12] and detected 9,053 trios with trans-associations at the 90% posterior probability threshold, each trio consisting of a cis-eQTL set, a cis-gene transcript, and a trans-gene transcript with evidence of association from the eQTL set on the trans-gene. We attempted to replicate our findings using data from two other studies of different tissue types – whole blood samples from the eQTLGen consortium [6] and dorsolateral prefrontal cortex samples from the CommonMind Consortium (CMC) [13].

**Figure 1.**
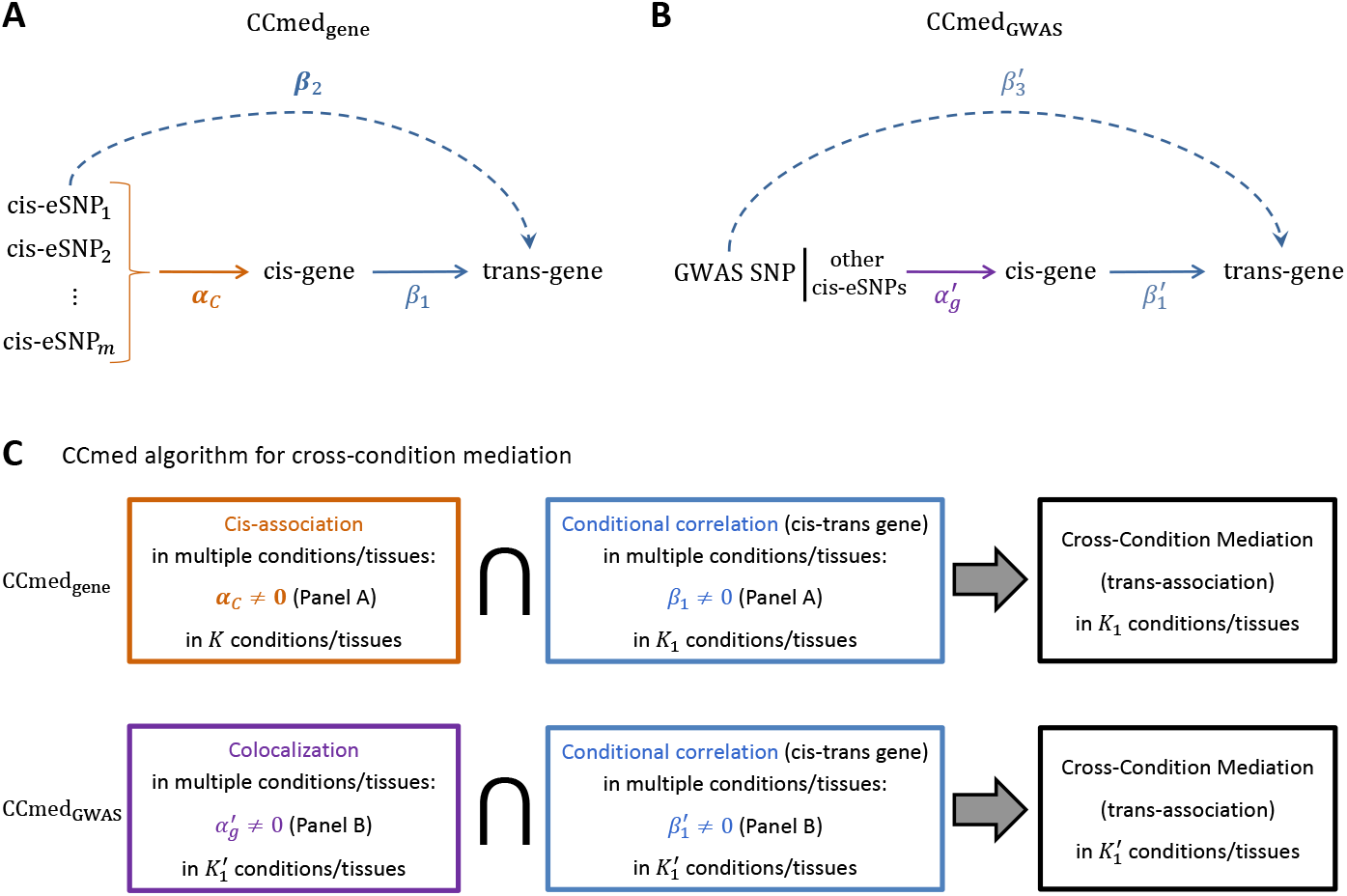
Two CCmed algorithms for identifying cis-mediated trans-associations across multiple conditions at the gene-level and for GWAS SNPs, respectively. (A) An illustration of the CCmed_gene_ model for mapping gene-level mediation and trans-association in a single tissue type. (B) An illustration of the CCmed_GWAS_ model for identifying trans-genes of a GWAS SNP mediated by cis-gene expression in a single tissue type. (C) A flowchart of the two CCmed algorithms to establish cross-condition (here, cross-tissue) trans-associations mediated by cis-gene expression levels.

In addition to mapping robust gene-level trans-associations in the genome, another major focus of this work is to study the trans-genes of GWAS SNPs and further understand how genetic variation affects complex traits – a long-standing question in genetics research. Increasing evidence has suggested that many susceptibility loci may affect complex traits/diseases via the modulation of their cisgene expression levels [14, 15, 16, 17]. Those cis-gene expression levels may further affect other distal genes in functional pathways, some of which also contribute to phenotypic variation. Here, by focusing on trait-associated SNPs identified from GWAS, we adapted the CCmed framework to identify trans-genes for GWAS SNPs, and proposed a CCmed_GWAS_ method (see Figure 1B&C). Different than CCmed_gene_ for gene-level trans-association, CCmed_GWAS_ focuses on one GWAS SNP at a time and aims to detect trans-genes for a GWAS SNP mediated by cis-gene expression while accounting for other cis-eQTLs in the gene region, i.e., a GWAS SNP |_other cis-eQTLs_ → cis-gene → trans-gene. CCmed_GWAS_ quantifies the joint probabilities of two conditions being simultaneously satisfied in at least some tissue types. The two conditions are: 1) the GWAS SNP is also a cis-eQTL conditioning on other cis-eQTLs; and 2) there exists non-zero conditional correlations between cis- and trans-expression levels conditioning on eQTL genotypes. As a proof-of-concept, we applied the CCmed_GWAS_ method to the multitissue eQTL data from 13 brain tissue types of the GTEx project, and detected suspected trans-genes for known schizophrenia (scz)-associated loci [18].

Not all of the trans-genes associated with disease susceptibility loci will be involved in the disease etiology. Considering the pleiotropic effects of many genetic variants [19, 20] and the potential false positive findings due to violations of assumptions required for mediation and association analyses, it is possible that only some proportion of the identified suspected trans-genes for GWAS loci are truly affecting disease risk. In order to further identify and validate the effects of the suspected trans-genes for 108 scz risk loci, we conducted three validation analyses. Among the suspected trans-genes for scz GWAS SNPs identified by CCmed_GWAS_, many of them show evidence of scz-risk associations in the validation analyses.

Many cis-mediated trans-associations have only weak effects, and will be undetectable via direct trans-association tests. In this work, we developed two tailored methods for detecting cis-mediated trans-associations across multiple conditions. A major innovation of our work is that we focused on cis-mediated trans-association effects, both at the gene-level and for GWAS-SNPs, and showed that those effects are not only prevalent in the genome, but also are often shared across related tissue types and are replicable across studies. The proposed CCmed framework takes only summary statistics as input and can be applied across different tissues/datasets/studies.

## Results

### Methods overview

In this work, we developed two cross-condition mediation analysis methods, CCmed_gene_ and CCmed_GWAS_, for detecting cis-mediated trans-associations at the gene-level and for GWAS SNPs, respectively. CCmed takes as input summary statistics from multiple studies/tissue-types/conditions, aiming to detect robust mediation and trans-association effects shared across conditions. Additionally, to validate the trait-associations of the identified suspected trans-genes for GWAS SNPs, we developed a two-sample Mendelian Randomization (MR) method ROBust to correlated and some INvalid instruments, termed “MR-Robin”. While the CCmed methods can be applied to many cross-condition settings (e.g. cell types), in this work we applied the methods to study cross-tissue mediation effects. For clarity and consistency between descriptions of the methods and data applications, we used the term “tissue” in lieu of “condition” in describing the methods.

#### CCmed_gene_ for mapping gene-level trans-associations mediated by cis-gene

CCmed_gene_ detects trios (eQTL set, cis-gene, trans-gene) showing evidence of crosstissue trans-association and mediation effects by quantifying the joint probability of two conditions being satisfied in at least *K*_1_ out of *K* tissue types. The two conditions are gene-level cis-associations and non-zero correlations between the expression levels of the cis- and trans-genes conditioning on the eQTL genotypes. Note that CCmed allows that the trios may also have a direct effect from the eQTL set to the trans-gene not completely mediated by the cis-gene expression (i.e., ***β***_2_ ≠ 0 in Figure 1A).

Specifically, for each trio (***L**_i_*, *C_i_*, *T_j_*), where ***L**_i_* is a set of eQTL genotypes for a cis-gene *i*, *C_i_* is the cis-gene expression level and *T_j_* is the expression level of a trans-gene *j*, we calculated the probability that *C_i_* mediates the effects of ***L**_i_* on *T_j_* in at least *K*_1_ tissue types, P_med,*ij*_, as follows:

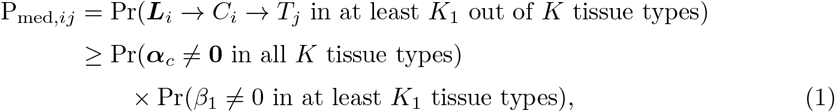

where ***α**_c_* is a vector of cis-association effects for the set of eQTLs in a single tissue type, and *β*_1_ is the conditional correlation of cis- and trans-gene expression levels in a single tissue type. We estimated ***α**_c_* and *β*_1_ for each tissue type and obtained the summary statistics. Note that here we omitted the subscript for tissue type to make notation simpler, though gene expression levels, cis-association and conditional correlation effects all vary by tissue type.

To quantify the cross-tissue cis-association probability, Pr(**α**_*c*_ ≠ **0** in all *K* tissue types) in (1), we first obtained the gene-level cis-association statistics for *M* cis-genes by *F*-tests. We applied a recently developed integrative association analysis approach, Primo [15, 21], to the association statistics to obtain the probabilities of 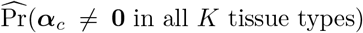 for gene *i* = 1, …, *M*. Similarly, by applying Primo to the conditional correlation statistics from *K* tissue types for the 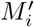 trans-genes for each cis-gene *i*, we obtained the probability of non-zero conditional correlation in at least *K*_1_ tissue types for all trans-genes of a cis-gene, 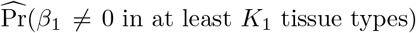. By taking the products of the two probabilities, we estimated a lower bound of the probability of gene-level cis-mediated trans-associations for each trio of (***L**_i_*, *C_i_*, *T_j_*) in at least *K*_1_ tissue types. Additional details are provided in the Methods section.

#### CCmed_GWAS_ for detecting trans-genes for GWAS SNPs

To detect trans-genes associated with GWAS SNPs of a complex trait via mediation analysis, CCmed_GWAS_ quantifies the joint probability of two conditions being satisfied in at least 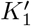 out of *K* tissue types. The two conditions are: (i) the GWAS SNP is also a cis-eQTL for the cis-gene conditioning on other cis-eQTLs, and (ii) there is a non-zero correlation between the cis- and trans-gene expression levels conditioning on the genotypes of eQTL and GWAS SNPs. Specifically, we calculate the probability that a GWAS SNP (*G_i_*) is affecting the expression of a distal gene (*T_j_*) in at least 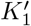 of *K* tissues via modulating its cis-gene expression (*C_i_*), while also allowing partial mediation, as follows:

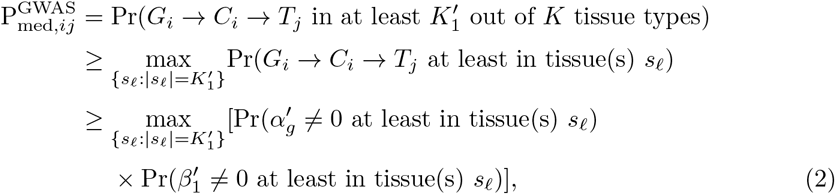

where *G_i_* is the genotype of a GWAS SNP of interest, *s*_*ℓ*_ is a set of tissue indices and is a subset of {1, 2, · · ·, *K*} with 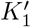 distinct tissue types. There are a total of 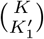 unique *s_ℓ_*’s. The parameter 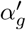 is the conditional cis-association statistic of the GWAS SNP of interest to a local gene’s expression conditioning on other lead eSNPs (an eSNP is an eQTL SNP, and a lead eSNP is the eSNP with the smallest eQTL *P*-value in the region), and a non-zero 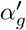 value implies the GWAS SNP is also an independent eQTL regulating the cis-gene expression *C_i_*. And 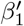 is the conditional correlation statistic of cis- and trans-gene expression levels conditioning on eQTL and GWAS SNPs. Both cis-association and conditional correlation statistics are calculated separately for each tissue type, and we again omitted the subscript for tissue type for simpler notation. We apply the Primo method to calculate the two probabilities in (2) to obtain a lower bound of the probability of trans-gene association of a GWAS SNP via cis-mediation.

CCmed_gene_ and CCmed_GWAS_ differ in the following major aspects: first, the cis-association statistic of CCmed_gene_ is calculated based on an *F*-test for a set of cis-eQTLs, whereas the cis-association statistic of CCmed_GWAS_ focuses on only the cis-association of a GWAS SNP, adjusting for other eQTLs. Second, in calculating the conditional correlation statistics, CCmed_gene_ adjusts for cis-eQTLs, and CCmed_GWAS_ also adjusts for the GWAS SNP in addition to cis-eQTLs. Third, it can be seen from our results that there are many robust gene-level cis-mediated trans-associations with effects shared across multiple tissue types, and we used *K*_1_ = 12 out of *K* = 13 brain tissues in detecting the more robust cross-brain-tissue trans-association analysis of the GTEx data. In contrast, the GWAS SNPs of a complex disease/trait may be eQTLs in only specific disease/trait-relevant tissue types and subsequently may have trans-associated genes in specific tissue types. As such, we suggest using a relatively small 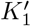 (and 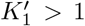 would ensure some robustness) for CCmed_GWAS_. In detecting the trans-genes for scz risk loci, we choose 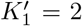 out of 13 brain tissue types in the analysis. Note that if a GWAS SNP has multiple cis-genes in the cis-region, we will separately consider them.

### Validating the trait-associations of the suspected trans-genes for GWAS SNPs

Not all of the trans-genes associated with GWAS SNPs for a complex disease/trait will be involved in the disease/trait etiology. Considering the pleiotropic effects of many genetic variants, only some of the trans-genes for GWAS SNPs are truly affecting disease risk or trait variation. Therefore, we conducted three validation analyses to assess the effects of suspected trans-genes of GWAS SNPs identified by CCmed_GWAS_ on the complex disease/trait of interest.

As shown in Figure 2, those validation analyses share a common rationale: considering a suspected trans-gene identified via CCmed_GWAS_, if the trans-gene is truly affecting the complex trait variation, then the local eQTLs of the trans-gene would also have associations to the complex trait; otherwise, they would not. Using an analogy, there is a known “ancestor” (GWAS SNP) and “descendant” (complex trait) relationship, and there are two suspected “fathers” (trans-genes) of the descendant. When the father information is not available (trans-gene expression not being measured in the GWAS study), then by examining the relatedness of the descendant (complex trait) to the local relatives from maternal side of the suspected fathers (local eQTLs for the suspected trans-genes) using only GWAS statistics, one can validate the paternal relationship (trait-association of the suspected trans-gene). Here we recapitalized on existing multitissue eQTL resources from the GTEx Project and the GWAS summary statistics from the Psychiatric Genomics Consortium (PGC), and conducted three types of validation analyses on the suspected trans-genes for scz loci detected from GTEx data via CCmed_GWAS_. We examined their risk-associations based on: 1) single-SNP GWAS summary statistics for local eQTLs of the suspected trans-genes; 2) a gene-based association test via transcriptome-wide association analysis (TWAS); and 3) a newly-developed two-sample Mendelian Randomization method – MR-Robin.

**Figure 2.**
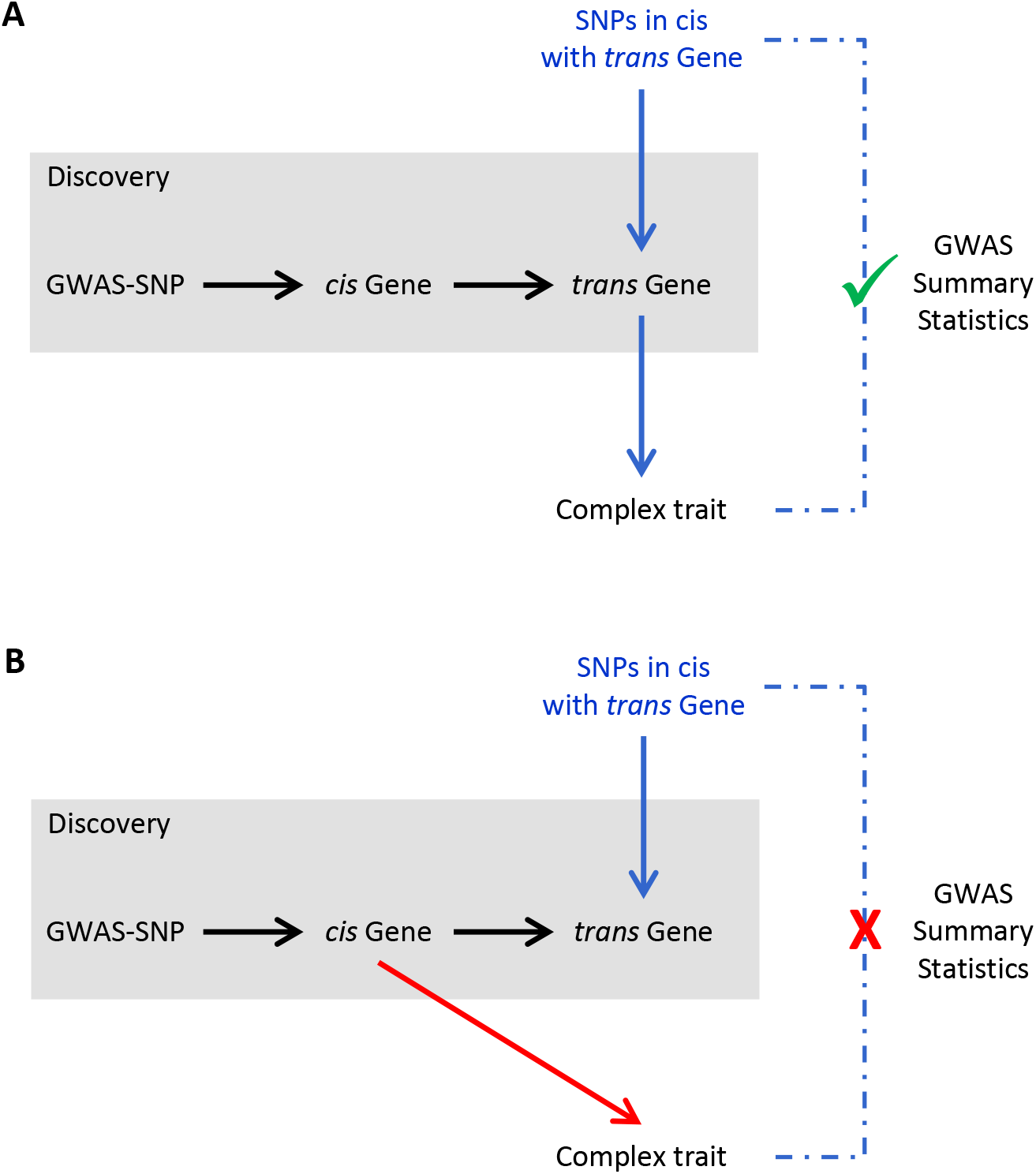
A conceptual illustration of the validation analyses to examine the trait-associations of the suspected trans-genes for GWAS SNPs. In the discovery analysis by CCmed_GWAS_, we identified the suspected trans-genes for GWAS SNPs. If a trans-gene is involved in the trait/disease etiology as in (A), we expect the SNPs in cis with the trans-gene to have higher than random association effects with the complex trait. When a trans-gene is not involved in the trait/disease etiology as in (B), we expect to observe the SNPs in cis with it not associated with the complex trait higher than random.

#### MR-Robin: a robust two-sample Mendelian Randomization method for assessing the causal effect of gene expression on a complex trait

MR-Robin assesses the causal effect of gene expression on a complex trait by integrating and recapitalizing on existing GWAS and multitissue eQTL statistics. Note that we applied MR-Robin in the validation analysis, though it can be generally applicable to assess the causal effect of a gene on a trait for discovery purposes.

Many existing methods have been proposed in the Mendelian Randomization literature to assess the causal effects of gene expression on a trait by harnessing Mendelian randomized genetic variants as instrumental variables (IVs) [22, 23, 24, 25]. A valid IV is a genetic variant that has effects on the complex trait completely mediated through gene expression [26], and is independent of unmeasured confounders of the mediator (gene expression) and outcome (complex trait). That is, there is no “horizontal pleiotropy” (a phenomenon that a genetic variant also affects the complex traits via other pathways not mediated through cis expression) [22] nor “correlated pleiotropy” [27] (a phenomenon that a genetic variant affect both mediator and outcome through a heritable shared factor, i.e. IVs are associated with the confounder). Note that valid IVs do not have to be the causal SNPs. The inclusion of invalid IVs in the MR analysis may lead to biased causal effect estimation and inference. A more detailed review of existing literature can be found in Methods section. In summary, most existing two-sample MR methods require a large number of IVs (here cis-eQTLs), and some methods require IVs to be nearly independent, limiting the applicability of those methods in assessing the causal effects of gene expression on complex trait.

The proposed MR-Robin method fills the gap. It requires only summary-level GWAS and multitissue eQTL statistics as input, considers multitissue eQTL effects for multiple IVs of a gene, allows IVs to be correlated and some of them to be invalid, and can be applied to genes with only a small number of cis-eQTLs. Specifically, traditional two-sample MR methods estimate the effect from gene to trait by taking the ratio of marginal effects of GWAS to eQTL associations, *β_yi_/β_xi_*, where *β_yi_* and *β_xi_* are the marginal GWAS and eQTL association effects, respectively, for SNP *i*. Due to linkage disequilibrium (LD) among SNPs and the pervasive horizontal and potential correlated pleiotropic effects in the genome [28, 27, 29], based on marginal summary statistics of one SNP *i* the estimated effect of gene to trait will be biased if any other SNP in the LD block has a horizontal or correlated pleiotropy effect. The magnitude of the bias depends on many factors including LD pattern, eQTL effects, and effects of horizontal or correlated pleiotropy. The bias is thus specific to each SNP (see Supplemental Materials for details).

MR-Robin considers the estimated effect of gene to trait from each IV as an observed value of the true effect plus a SNP-specific bias. By jointly considering multiple IVs, MR-Robin decomposes the estimated effects of multiple IVs into two components – a concordant effect shared across IVs and a discordant component allowing some IVs to be invalid with SNP-specific deviations from the true effect. MR-Robin makes the estimation identifiable by taking advantage of the multitissue eQTL effects for multiple IVs of a gene and treating them as the response variable in a reverse regression, with GWAS effect estimates as the predictor. The rich multitissue eQTL effect information in the response variable allows the estimation of SNP-specific random-slopes (i.e. deviated effects) due to potential invalid IVs. In contrast, to our knowledge, all existing MR methods use only single-tissue eQTL effects. Thus, with only a limited number of potentially correlated IVs, MR-Robin can accurately test the true effect from gene to trait by testing the shared fixed effect of correlation of eQTL and GWAS effects across IVs. A major innovation of MR-Robin is the use of rich information in multitissue eQTL effects in two-sample MR analyses. Additionally, correlated IVs are allowed, and LD among SNPs and estimation uncertainty are accounted for, with a *P*-value for each gene being calculated from resampling. A detailed description of the MR-Robin method can be found in the Methods section.

### Simulations to evaluate CCmed in detecting robust cis-mediated trans-associations across conditions

In this section, we evaluated the performance of CCmed_gene_ and CCmed_GWAS_ through simulation studies. We showed that when individual trans-eQTL effects are weak but are mediated by cis-gene expression with effects shared across tissue-types, the CCmed algorithms can borrow information across tissue types to improve power and detect cis-mediated trans-associations, while controlling the false discovery rate (FDR). We also compared with directly testing for trans-associations in each single tissue type and showed the advantages of the proposed algorithms. In each of the algorithm evaluations, we simulated 2.5 × 10^5^ trios (SNP(s), cis-gene expression, trans-gene expression) for 350 subjects in *K* = 10 tissues.

#### The performance of CCmed_gene_ in identifying robust gene-level trans-associations

We simulated 500 sets of cis-eQTLs for 500 cis-gene expression levels. For each set of cis-eQTLs, we simulated the genotypes of 10 correlated eSNPs with pairwise correlation of 0.3. Based on the genotypes, in each tissue type, we randomly selected 1 SNP as the causal eSNP to generate cis- and trans-gene expression levels. The causal eSNPs varied across tissues. For each pair of a cis-eQTL set and a cis-gene, we generated 500 trans-gene expression levels. There are a total of 250, 000 trios of (cis-eQTL, cis-gene, trans-gene). We generated cis- and trans-gene expression data from 10 correlated tissue types. Expression data was simulated such that cis-gene expression was associated with the cis-eQTL set in a varying proportion of the tissues (including none), and that non-zero conditional cis-trans gene expression correlation was present in a subset of the cis-trans gene pairs in a varying proportion of the tissues (including none). Among those trios with non-zero cis-mediated trans-associations, 50% of them also had non-zero direct effect from SNPs on the trans-gene expression levels. Effect sizes were simulated to mimic weak total trans-associations as observed in the GTEx study. See Supplemental Materials for additional simulation details. In the simulation studies, we are interested in detecting the trios with cis-association and conditional expression correlation in at least 9 out of 10 tissue types.

Table 1A presents the powers as well as the true and estimated FDRs to detect gene-level trans-associations mediated by cis-gene in at least *K*_1_ = 9 tissue types at a threshold of 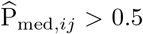, 0.8 and 0.9, respectively. As a comparison, we also obtained the *P*-values of *F*-statistics that directly test for the total gene-level trans-association effects. For each of the thresholds used in the CCmed_gene_ approach, we applied the corresponding estimated FDR to the 250, 000 by 10 matrix of *P*-values of direct tests and obtained the trios with significant total gene-level trans-associations in at least 9 tissues. The powers of the direct test are low as expected due to weak trans-association effects, stringent cross-tissue association criteria (i.e., 9 out of 10) and multiple testing burden. In contrast, CCmed_gene_ greatly improved the power by exploring mediation-based association and by borrowing information across tissues. The FDRs are well-controlled by both methods.

**Table 1.** Simulation results evaluating the performance of CCmed. estP := estimated probability; estFDR := estimated FDR.(A) Simulation to compare CCmed_gene_ with direct test for trans-association in detecting associations in at least *K*_1_ = 9 tissue types. (B) Simulation to compare CCmed_GWAS_ with direct test for trans-association in detecting associations in at least 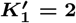 tissue types.

| (A) |  |  |  |  |  |  |  |  |  |
| --- | --- | --- | --- | --- | --- | --- | --- | --- | --- |
| Method | Association in at least $K_1 = 9$ tissue types | | | | | | | | |
| | $\text{estP} \geq 0.50$ | | | $\text{estP} \geq 0.80$ | | | $\text{estP} \geq 0.90$ | | |
|  | Power (%) | true FDR(%) | estFDR (%) | Power (%) | true FDR(%) | estFDR (%) | Power (%) | true FDR(%) | estFDR (%) |
| CCmed<sub>gene</sub> | 68.0 | 3.2 | 7.7 | 60.2 | 2.2 | 4.0 | 54.7 | 2.2 | 3.1 |
| Direct test | 7.7% estFDR |  |  | 4.0% estFDR |  |  | 3.1% estFDR |  |  |
|  | 31.6 | 9.4 | - | 25.9 | 4.8 | - | 23.9 | 3.6 | - |

| (B) |  |  |  |  |  |  |  |  |  |
| --- | --- | --- | --- | --- | --- | --- | --- | --- | --- |
| Method | Association in at least $K'_1 = 2$ tissue types | | | | | | | | |
| | $\text{estP} \geq 0.50$ | | | $\text{estP} \geq 0.80$ | | | $\text{estP} \geq 0.90$ | | |
|  | Power (%) | true FDR(%) | estFDR (%) | Power (%) | true FDR(%) | estFDR (%) | Power (%) | true FDR(%) | estFDR (%) |
| CCmed<sub>GWAS</sub> | 83.2 | 2.3 | 2.9 | 79.2 | 0.9 | 0.9 | 76.7 | 0.6 | 0.4 |
| Direct test | 2.9% estFDR |  |  | 0.9% estFDR |  |  | 0.4% estFDR |  |  |
|  | 60.6 | 0.8 | - | 55.4 | 0.2 | - | 52.3 | 0.1 | - |

#### The performance of CCmed_GWAS_ in identifying cis-mediated trans-genes for one (GWAS) SNP in selected tissue-types

In this simulation, we simulated cis-gene expression levels being affected by 3 correlated eQTLs with correlation 0.3. We focused on one of them as the (GWAS) SNP of interest and generated the trans-gene expression levels being affected by the SNP in selected tissue types. Effects sizes were simulated to mimic scenarios with weak to moderate effects in selected and limited tissue types. See Supplemental Materials for additional simulation details.

Table 1B compares the performance of CCmed_GWAS_ with the direct test for the total trans-association effects for the (GWAS) SNP of interest to identify associations in at least 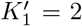 tissues. With stronger effects on average and smaller 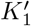 than those in the previous simulation, the direct test enjoys reasonable power this time. However, with CCmed_GWAS_, the power can still be improved by around 40% to 50% in this simulation setup with the same FDR control.

### Simulations to evaluate the performance of MR-Robin when IVs are correlated, some being invalid, and/or limited in number

In this section, we conducted simulation studies to compare MR-Robin with competing models and methods. We showed that MR-Robin is robust to the inclusion of correlated and some proportions of invalid IVs even when the number of IVs is small. We compared MR-Robin to three competing models and several competing methods in the literature. See Supplemental Materials for details of the simulation and the description of competing models. The first model is a single-tissue MR model with no intercept.The second and third models both use multitissue eQTL effect estimates, either as the response variable or predictor in the regression but do not consider SNP-specific deviation effects (random-slopes), unlike MR-Robin. We also compared MR-Robin to three existing MR methods in the literature that are based on summary statistics and are robust to invalid IVs: MR-RAPS [30], MR-Egger [31], and MRMix [29].Note that those three methods were developed for settings where many independent genetic variants are available as candidate IVs, for example when analyzing a polygenic trait as a mediator for other complex traits. Therefore, they may not be suited for our target settings in which gene expression (with only a limited number of correlated eQTLs/IVs) is considered as the mediator. Most of those existing methods also do not allow the IVs to be correlated. And they are all developed for single-tissue eQTLs. Nonetheless, we included the methods for comparison.

Table 2 shows the type I error rate and power comparison in the presence of 0, 10, …, 70% invalid IVs, allowing IVs to be correlated (pairwise LD *r*^2^ < 0.5) over 10,000 simulations of 10 LD blocks with 20 correlated SNPs in each block, before the IV selection (details in Supplemental Materials). As shown in the table, whereas competing methods are unable to control the type I error rate when there are any invalid instruments and instruments are in LD, MR-Robin maintains reasonable control of the type I error rate if a majority of instruments are valid (up to 30% invalid IVs). The last three methods in the table were developed for independent instruments; since they do not account for correlation (LD) among the instruments, they do not control the type I error rate even when all instruments are valid. In Supplemental Materials, Tables S1-3, we compared the type I error rates and powers using alternative LD selection criteria for the IVs (pairwise LD *r*^2^ < 0.8, 0.3 or 0.01). Moreover, in Supplementary Table S4, we showed that even when the number of available IVs is very small (3-10), the proposed MR-Robin can still yield reasonable results if the small number of IVs are relatively less dependent. Last but not least, we want to emphasize that when IVs are correlated, if one IV is an invalid IV, all the other correlated IVs are also affected to some degree, and as such the random-slope model of MR-Robin fits the need for allowing correlated IVs when considering the effect of gene expression on complex trait.

**Table 2.** Simulation results evaluating the performance of MR-Robin. Averaged type I error rates and power over 10,000 simulations are shown by percentage of valid instruments. 10 LD blocks were simulated, with one true eQTL per LD block. Instruments were selected sequentially: the eSNP with the strongest association with gene expression was selected, and the next selected eSNP is the strongest-associated SNP remaining also with LD *r*^2^ < 0.5 with any already-selected eSNPs.

| Method | Proportion of valid IV (%) |  |  |  |  |  |  |
| --- | --- | --- | --- | --- | --- | --- | --- |
|  | 100 | 90 | 80 | 70 | 60 | 50 | 30 |
|  | Type I error rate |  |  |  |  |  |  |
| MR-Robin | 0.046 | 0.058 | 0.063 | 0.078 | 0.094 | 0.110 | 0.165 |
| A single tissue MR model with no intercept | 0.310 | 0.365 | 0.395 | 0.413 | 0.432 | 0.451 | 0.446 |
| A multitissue MR model with a fixed slope and no intercept | 0.047 | 0.075 | 0.094 | 0.109 | 0.124 | 0.138 | 0.145 |
| Random Intercept | 0.048 | 0.071 | 0.093 | 0.107 | 0.122 | 0.133 | 0.141 |
| MR-RAPS | 0.286 | 0.629 | 0.801 | 0.875 | 0.908 | 0.919 | 0.913 |
| MR-Egger | 0.141 | 0.219 | 0.263 | 0.302 | 0.325 | 0.332 | 0.343 |
| MRMix | 0.154 | 0.218 | 0.274 | 0.329 | 0.374 | 0.426 | 0.494 |
|  | Power |  |  |  |  |  |  |
| MR-Robin | 0.978 | 0.932 | 0.880 | 0.824 | 0.768 | 0.720 | 0.589 |
| A single tissue MR model with no intercept | 0.995 | 0.969 | 0.937 | 0.903 | 0.878 | 0.851 | 0.803 |
| A multitissue MR model with a fixed slope and no intercept | 0.999 | 0.948 | 0.885 | 0.822 | 0.766 | 0.713 | 0.614 |
| Random Intercept | 0.999 | 0.949 | 0.886 | 0.824 | 0.770 | 0.718 | 0.609 |
| MR-RAPS | 1.000 | 0.995 | 0.992 | 0.989 | 0.982 | 0.976 | 0.966 |
| MR-Egger | 0.888 | 0.805 | 0.739 | 0.675 | 0.635 | 0.614 | 0.537 |
| MRMix | 0.527 | 0.527 | 0.523 | 0.516 | 0.530 | 0.540 | 0.545 |
|  | Avg number of SNPs selected (valid/invalid) |  |  |  |  |  |  |
| All methods | 35.2/0 | 31.5/3.5 | 28.0/7.0 | 24.5/10.5 | 20.9/14.2 | 17.4/17.6 | 10.4/24.6 |

### Mapping cross-tissue gene-level trans-associations mediated via cis-gene expression levels in the 13 GTEx brain tissue types

We applied CCmed_gene_ to data from the 13 brain tissue types of the GTEx project to identify gene-level cis-mediated trans-associations in the brain tissues. There are 5,347 autosomal genes with a posterior probability > 90% of gene-level cis-association in all 13 brain tissues. For each of these 5,347 cis-genes, we further assess the conditional correlation to each of the trans-gene expression levels in each brain tissue type, and subsequently calculate the cross-tissue probability of mediation. At a threshold of 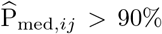, see equation (1), we identified a total of 9,053 trios (cis-eSNP set, cis-gene expression, trans-gene expression) showing evidence of gene-level cis-mediated trans-associations in at least 12 out of 13 brain tissue types (with an estimated FDR = 2.0% at 90% posterior probability). These trios included 604 unique cis- and 2509 unique trans-genes.

To replicate the trans-associations identified by CCmed_gene_, we used data from the eQTLGen and the CommonMind Consortia (CMC) as two replication studies [6, 13]. The eQTLGen Consortium, which focused on the 10,317 trait-associated SNPs, performed a meta-analysis of trans-eQTL association statistics based on whole blood samples of 31,684 individuals from 37 datasets [6], and has provided SNP-level blood-tissue trans-association *P*-values for those trait-associated SNPs. To replicate our gene-level trans-association findings with eQTLGen results, for each (*C_i_*,*T_j_*) cis-trans pair in a CCmed_gene_ mediation trio, we obtained the eQTLGen trans-association *P*-values between expression of *T_j_* and each SNP in cis with *C_i_*. As a comparison, we also obtained the eQTLGen trans-association *P*-values for randomly selected cis-trans gene pairs. In the QQ-plots of Figure 3A, the eQTLGen trans-association *P*-values for cis-SNPs and trans-genes identified by CCmed_gene_ (cyan points) show a much stronger enrichment of association than randomly selected sets (black points). This enrichment is present despite the discovery and replication analyses using different tissue types, brain versus blood, respectively, suggesting that CCmed_gene_ identifies cis-gene-mediated trans-associations that are robust across tissue types. Note that since the eQTLGen consortium focuses on only the trait-associated SNPs (which are enriched for cis-/trans-eQTLs), and moreover the sample size of eQTLGen is very large, the eQTLGen results for randomly selected cis-trans gene pairs are also enriched for trans-associations.

**Figure 3.**
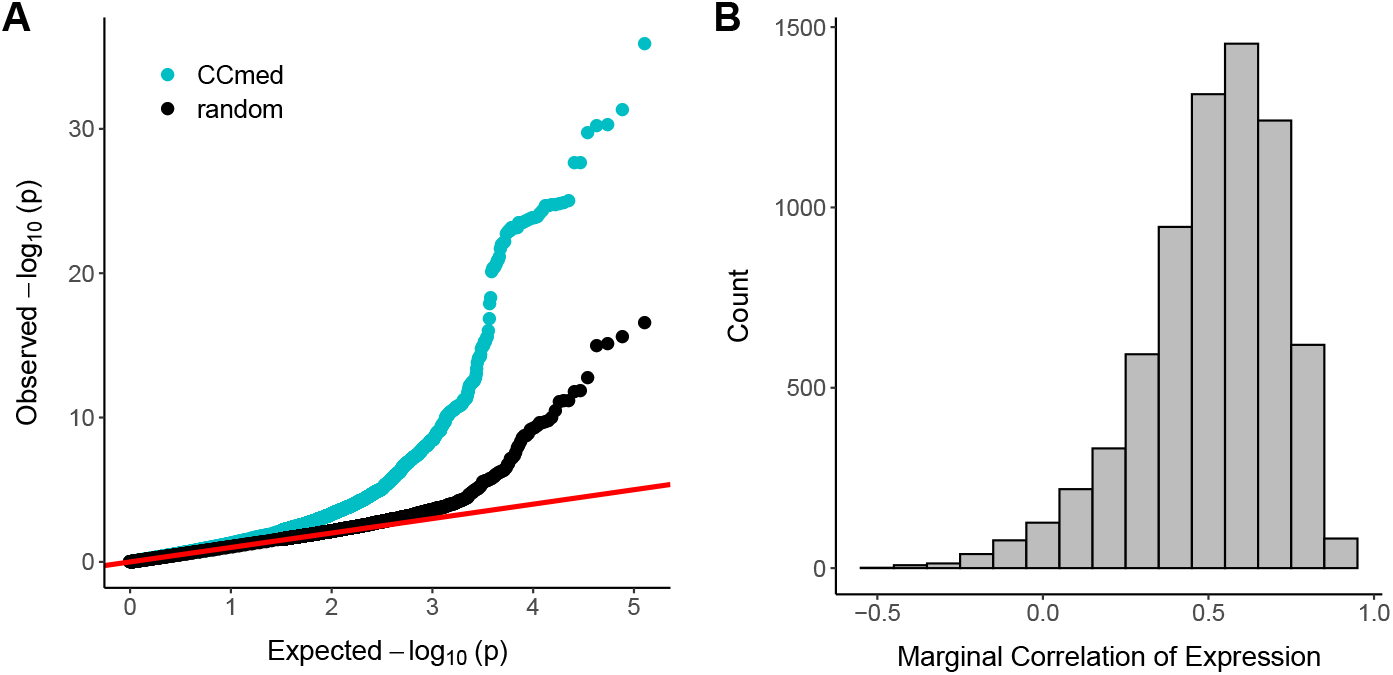
CCmed_gene_ trans-associations are replicated in other datasets. (A) QQ-plot of −log_10_(P) comparing trans-associations identified by CCmed_gene_ (cyan points) to randomly selected trans-associations (black points) in the eQTLGen blood tissue trans-eQTL study of trait-associated variants. (B) Marginal correlations in CMC Dorsolateral Prefrontal Cortex samples of cis-trans pairs in mediation trios identified by CCmed_gene_.

Next, we examined the cis-trans gene-gene correlations for gene pairs identified by CCmed_*gene*_ using data from the CMC [13]. CMC has generated DNA and RNA sequencing data from postmortem brain samples from donors with schizophrenia and bipolar disorder, and from subjects with no neuropsychiatric disorders (see Supplemental Materials for detailed data descriptions). For the cis-trans gene pairs identified by CCmed_gene_ using the GTEx data, Figure 3B shows their marginal expression correlations in the CMC data. As shown by the histogram, most cistrans pairs in mediation trios identified by CCmed_gene_ have moderately to strongly correlated expression levels in the dorsolateral prefontal cortex samples of CMC. Since 80% (FDR≤ 5%) of the genes in the CMC data are reported to have at least one cis-eQTL [13], the presence of cis-association and correlation of cis-trans gene expression implies gene-level trans-association and mediation effects being also present in the dorsolateral prefrontal cortex tissue. That is, we observed evidence of replication in the CMC data for our trans-associations findings from GTEx data identified by CCmed_*gene*_.

With the discovery analysis using GTEx data and replication analyses using eQTLGen and CMC data, we showed that cis-mediated trans-associations are abundant in the genome and a substantial proportion of them can be replicated across studies, tissue types, and populations (e.g. healthy individuals from GTEx versus diseased individuals in CMC). This is not surprising because each tissue type is a mixture of many cell types, and both cis-association and gene expression correlation may have effects shared across many cell types. The detection and availability of robust trans-eQTLs can be further used in many related areas, for example, in improving the expression imputation in TWAS analysis, or in constructing gene networks. Cross-condition mediation analysis, as a complementary approach to the direct trans-association test, will help to integrate data from multiple studies and to detect moderate-to-weak mediation and trans-association effects.

### Detecting trans-genes associated with 108 known schizophrenia loci in GTEx brain tissues

To detect trans-genes whose expression levels may be associated with schizophrenia risk, we applied CCmed_GWAS_ to the 108 scz susceptibility loci reported by the PGC consortium [18]. Of the 128 reported GWAS SNPs in these loci, 103 were genotyped in the GTEx data, 21 were captured by a SNP in strong LD (for lead GWAS SNPs in the region, we substituted the most significant SNP in the gene region present in GTEx; and for secondary GWAS SNPs, we substituted the nearest SNP mapped in GTEx that reached genome-wide significance), 1 SNP was not present nor captured by a SNP in high LD, and 3 non-autosomal SNPs were excluded in the analysis. For each of the 124 scz-associated SNPs we analyzed, there were multiple cis-genes in the cis-region. And there were a total of 1,643 unique cis-genes we considered for those 124 GWAS SNPs, with all cis-genes being expressed in all brain tissues and having at least 1 brain eQTL reported by GTEx.

For each (GWAS SNP, cis-gene) pair, we conducted regression (6) described in the Methods, regressing cis-gene expression on the GWAS SNP genotype and adjusting for GTEx-reported eQTLs, to obtain the cis-association statistics in each tissue type. We applied Primo [15] to the *M_g_* × *K* matrix of cis-association statistics {*F*^G^} where *M_g_* is the number of (GWAS SNP, cis-gene) pairs, and estimated the probability of each GWAS SNP being also an eQTL in at least 2 tissue types, conditioning on other cis-eQTLs. There were 40 (GWAS SNP, cis-gene) pairs showing evidence of cis-association in at least 2 tissue types (with cross-2-tissue cis-association posterior probability > 80%).

For each of the 40 (GWAS SNP, cis-gene) pairs, we further conducted conditional correlation analysis with each of its trans-genes in each tissue type using the regression model (7) described in Methods to obtain a matrix of conditional correlation statistics, 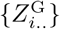, conditioning on cis-eQTL and GWAS SNP genotypes. We applied Primo [15] to 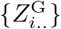 to estimate the probability of non-zero conditional cis-trans gene-expression correlation for each possible pair of tissues *s_ℓ_*. Using equation (2), we obtained a lower bound estimate of the probability of cis-mediated trans-association in at least two brain tissue types for each trio, and identified 1492 (GWAS SNP, cis-gene, trans-gene) trios with 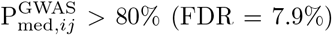 with 1418 unique trans-genes.

### Validating schizophrenia-risk-associations of suspected trans-genes based on GWAS summary statistics from PGC

To examine whether the 1418 identified trans-genes for the scz-GWAS SNPs are truly associated with scz risk, we conducted a series of validation analyses on those suspected genes. Those validation analyses are based on a common rationale (as shown in Figure 2). Considering a suspected trans-gene identified via CCmed_GWAS_, if the trans-gene is truly associated with the complex trait of interest (here, schizophrenia), then the local eQTLs of the trans gene would also have associations to the complex trait; otherwise, they would not. Therefore, by examining the trait-association statistics from existing GWAS for SNPs in cis with the suspected trans-genes, one could distinguish the trans-genes truly associated with complex trait versus those ones that may be only co-expressed with the scz-associated cis-genes. Note that suspected trans-genes not having any local eQTLs cannot be checked nor validated by those analyses.

First, we examined the single-SNP scz GWAS summary statistics from PGC for local eQTLs of suspected trans-genes. For each suspected trans-gene identified from GTEx via CCmed_GWAS_, we checked the GWAS scz-association *P*-values for the local eQTLs of the suspected trans-gene. The local eQTLs are eQTLs reported in at least one GTEx brain tissue reported by the GTEx consortium [12]. There are 1158 out of 1418 suspected trans-genes having at least one GTEx reported local eQTL that could also be mapped to the PGC GWAS summary statistics. We checked the scz-associations of the local SNPs of those genes and found that 589 genes (50.9%) had at least one local eQTL with GWAS scz-risk-association *P*-value < 0.05. Figure 4A shows the histogram of PGC GWAS *P*-values for the local eQTLs of the 1158 suspected trans-genes. Based on the histogram, the local eQTLs for suspected trans-genes identified by CCmed_GWAS_ are highly enriched for associations with schizophrenia-risk.

**Figure 4.**
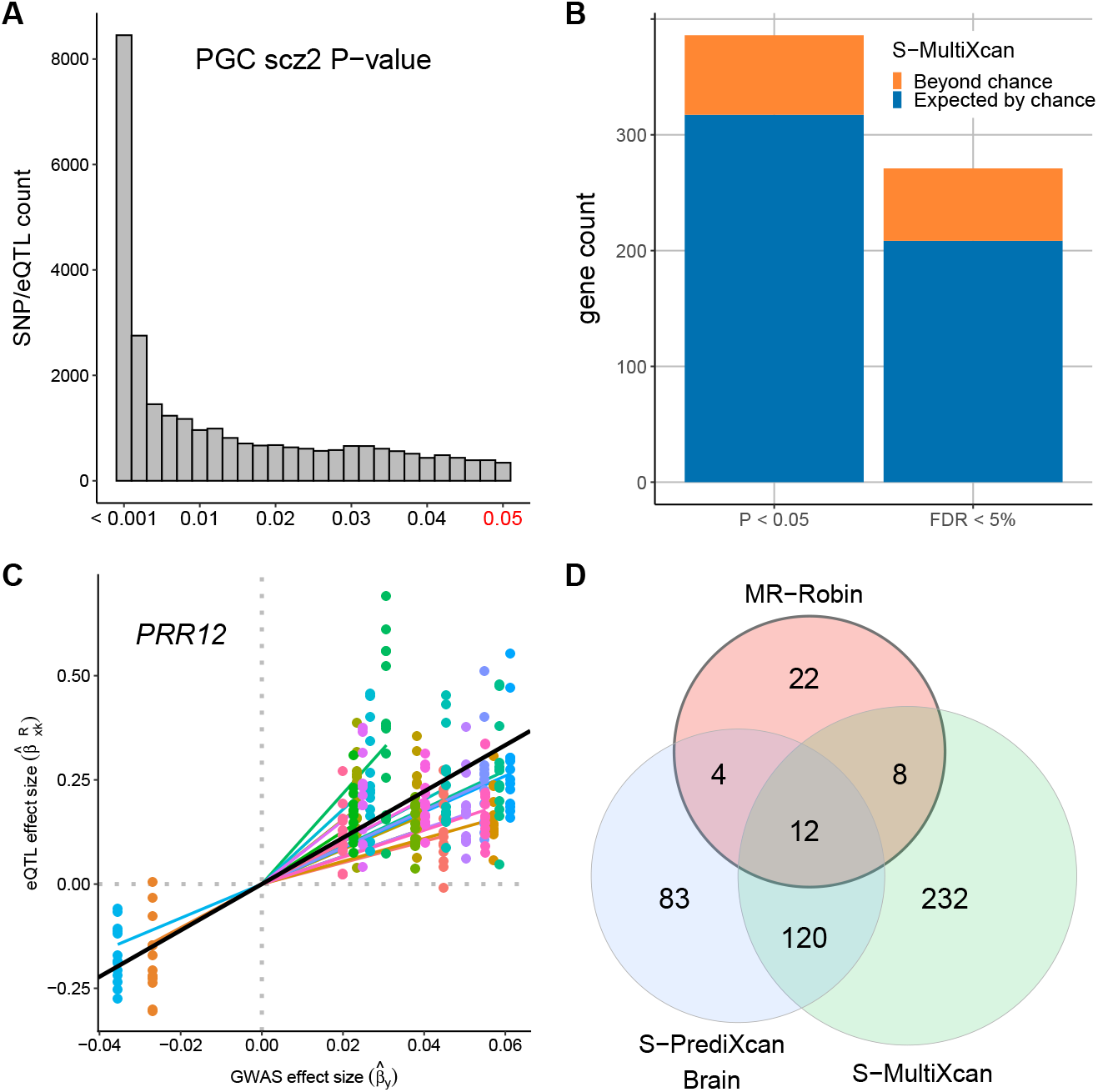
Validation of CCmed_GWAS_ trans-associations. (A) Among the 1418 trans-genes for 124 scz-SNPs detected from GTEx data via CCmed_GWAS_, 1158 trans-genes have at least 1 GTEx reported cis-eQTL, with a total of 124,619 eQTLs also genotyped in the GWAS data. Here we plot the GWAS scz-risk-association P-values for those 124,619 eQTLs (and zoomed-in to focus on those with GWAS *P* < 0.05). (B) Counts of suspected trans-genes showing evidence of schizophrenia-risk association based on multitissue TWAS analysis. Colors correspond to counts that would be expected by chance using randomly selected genes (blue) and the counts beyond what would be expected by chance (orange) among CCmed-identified suspected trans-genes. (C) An example of a suspected trans-gene validated by MR-Robin (p = 0.001). The eQTL effect size estimates (y-axis) are plotted against GWAS effect size estimates (x-axis) for SNPs used in the analysis. Points are colored by SNP. Colored lines represent SNP-specific slope estimates. The slope of the black line is the fixed effect estimate from the MR-Robin reverse regression, and implies a non-zero mediation effect of the gene *PRR12* on scz. (D) Venn diagram of gene counts of MR-Robin, brain-tissue-based S-PrediXcan and S-MultiXcan results at the cutoff of *P* < 0.05.

Next, we obtained the TWAS *P*-values for scz-risk associations for all genes from a recent analysis by Barbeira, *et al.* [32]. The TWAS analysis was conducted using PGC scz-risk GWAS summary statistics [18] and predicted transcriptome data with multitissue eQTL data from GTEx (V6p) as reference panels. We examined the results from S-MultiXcan, which used a multitissue eQTL reference panel of 44 GTEx tissue types (including brain and non-brain tissue types). The barplot in Figure 4B summarizes the gene counts in the predicted transcriptome dataset at the cutoffs of *P*-value < 0.05 and FDR < 5%. We compared the counts of the 1418 trans-genes identified by CCmed_GWAS_ present at each cutoff (total bar height) to the average counts of 1418 randomly selected genes present at each cutoff (blue bar), averaged across 100 randomly selected sets. Out of 1290 CCmed_GWAS_ transgenes tested in S-MultiXcan, 386 (29.9%) and 271 (21.0%) genes were significant at *P* < 0.05 and FDR < 5%, respectively. The suspected trans-genes identified by CCmed_GWAS_ have higher than random gene-level scz-risk-associations based on the results from Barbeira, *et al.* [32].

Both single-SNP and TWAS-based gene-level analyses showed the enrichment of scz-risk associations for the suspected trans-genes identified by CCmed_GWAS_ from GTEx data. To precisely identify the trans-genes causal for schizophrenia risk, we applied the proposed two-sample MR method, MR-Robin, to the identified suspected trans-genes. Note that only trans-genes with multiple local cross-tissue eQTLs can be properly analyzed using MR-Robin. If the gene has only 1 cis-eQTL, MR-Robin is reduced to a single-IV analysis, which can be heavily affected by the validity of the IV with assumptions that cannot be adequately checked in general. Therefore, in the following analysis, we restricted the MR-Robin analysis to the suspected trans-genes with at least three local eQTLs (within 1 Mb of transcription start site) that have median eQTL *P*-value being < 0.05 across 13 GTEx brain tissues. We formed the set of IVs by iteratively selecting the SNP with the smallest median eQTL *P*-value and having pairwise LD *r*^2^ < 0.5 with each SNP already selected. Among the 1418 identified suspected trans-genes, 493 had at least 3 such SNPs selected (median *P* < 0.05 and pairwise LD *r*^2^ < 0.5) and were tested using MR-Robin. At the *P*-value cutoff of 0.05 based on MR-Robin, 46 of the trans-genes showed significant dependence of GWAS and eQTL statistics, implying significant scz-risk associations for the trans-genes (see Supplementary Materials for details of the 46 genes validated by MR-Robin). The scatterplot in Figure 4C shows an example of a significant gene (*PRR12*, with MR-Robin *P* = 0.001). The eQTL effect sizes in the GTEx brain tissues were plotted against the GWAS effects sizes in the PGC dataset for the selected SNPs for the gene (*PRR12*). Despite some SNPs having potentially larger deviation from the others – indicated by the random slopes (colored lines) deviating from the fixed effect estimate (black line) – the plot shows a clear pattern of association between the magnitude of eQTL effects and magnitude of GWAS effects, implying that the expression levels of the gene affects scz risks. In Figure 4D, we examined the overlaps of results based on MR-Robin and TWAS. In addition to the S-MultiXcan TWAS results, we also examined overlap with single tissue S-PrediXcan TWAS results using GTEx Brain tissues, summarized by taking the minimum *P*-value across 10 GTEx V6p Brain tissues. Among the 46 causal trans-genes identified by MR-Robin, 24 are also implied by either S-PrediXcan Brain or S-MultiXcan. Among the 22 genes that are implied by neither TWAS method, 17 were included in two or fewer brain tissues in the TWAS results, likely due to poor expression prediction; and, the genes *DGCR6* and *MDGA2* have been reported in the literature with evidence of associations to schizophrenia [33, 34, 13].

There is significant interest in identifying the mechanisms through which GWAS SNPs affect complex traits. Much research examining the effects of GWAS SNPs on gene expression has focused on cis-associations. Here we introduced a mediation method to identify trans-associations of GWAS-SNPs and proposed a series of validation analyses to identify which of the suspected trans-genes may be associated with complex trait variation such as disease risk. As a proof-of-concept, we analyzed scz-risk associated SNPs and detected their trans-genes based on CCmed_GWAS_. With validation analyses, we showed that the detected trans-genes are enriched with scz-risk-associations. A highlight of the validation analysis is that using a newly proposed two-sample MR method, MR-Robin, we leveraged the rich information provided by the multitissue eQTL statistics from GTEx and the GWAS statistics from PGC, and further fine-mapped 46 trans-genes with potential causal associations to scz risk, many of which are also supported by other evidence.

## Discussion

In this work, we focused on studying trans-associations mediated by cis-gene expression levels. The cis-mediated trans-association effects have a direct mechanistic interpretation; moreover, we show that many of those trans-associations may have effects shared across functionally related tissue types, cell types, studies or cellular conditions, and are robust across conditions and replicable. To detect trans-associations via cross-condition mediation analyses at the gene level and for GWAS SNPs, we proposed two methods, CCmed_gene_ and CCmed_GWAS_, respectively. Both methods take as input the summary statistics of cis-eQTL-to-expression association and conditional cis-trans expression correlations from multiple tissue-types, cell-types, studies or cellular conditions, and quantify the probability of cis-mediated trans-associations in multiple conditions. By applying CCmed_gene_ to data from the 13 brain tissues of the GTEx project, we identified cis-trans gene pairs with gene-level trans-association effects in most brain tissues. Our findings are replicable with many cis-trans gene pairs showing evidence of trans-association effects in two different tissue types from two other studies. As a proof-of-concept for CCmed_GWAS_, we applied the method to 108 schizophrenia susceptibility loci and identified the trans-genes for scz GWAS SNPs with cis-mediated trans-association effects in at least 2 out of 13 GTEx brain tissues. By harnessing GWAS and GTEx multitissue eQTL statistics for SNPs in cis with the identified suspected trans-genes, we further conducted validation analyses to examine the trait-associations of suspected trans-genes using single-SNP GWAS statistics, TWAS-based analyses, and a proposed two-sample MR approach (MR-Robin).

Trans-acting genetic effects on distal gene expression are ubiquitous in the genome; however, they are often found to act in a tissue-specific or cell-type-specific manner. Using standard trans-association tests correlating SNP genotypes with each distal gene expression in the genome and adjusting for multiple comparison, the detection and replication of trans-association effects require large sample sizes. Our work describes a complementary approach for studying trans-associations mediated by cis-gene expression, one that can increase power to detect robust trans-associations by borrowing information across studies/tissue-types/conditions. By analyzing multitissue eQTL data from GTEx and replicating our findings, we showed that many cis-mediated trans-associations are robust and replicable in different studies. Here we applied the methods to data from multiple brain tissues, while the proposed methods and analyses can be used and generalized to recapitalize on existing eQTL data from many different tissue types and studies. Given that trans-eQTLs account for a substantial amount of expression heritability in the genome, we anticipate that replicable trans-association findings would provide new opportunities in advancing related research areas in integrative analyses involving eQTL data – for example, in improving expression imputation in TWAS analysis. Furthermore, the proposed trans-association analysis may be able to identify convergent gene networks when multiple cis-genes mediate a common trans-gene, or help infer master regulators of disease-relevant gene networks [35].

Another innovation of this work is the development of MR-Robin – a two-sample MR approach to assess the causal effects of gene expression on a complex trait of interest. The method is used as a validation approach in the work but can be applied as a discovery method in other contexts. Compared to existing two-sample MR methods, a major advantage of MR-Robin is the use of multitissue eQTL summary statistics. Selecting the eQTLs with cross-tissue effects will improve the reproducibility of eQTL effects and subsequent findings across two samples - the eQTL reference and the GWAS data. More importantly, MR-Robin is based on a mixed-effect model with multitissue eQTL effects from multiple IVs as the response variable and GWAS effects as the predictor, testing for non-zero correlation of eQTL and GWAS effects from multiple IVs. The rich information in multitissue eQTL effects allow the estimation of a shared fixed-effect correlation of eQTL and GWAS effects across all IVs, as well as the estimation of SNP-specific deviation due to invalid IVs captured by the random-slope. In contrast, existing models and methods based on the deconvolution of mixture distributions or penalized regressions in general require a large number of IVs to achieve stability in estimation. With both simulation studies and real data analyses, we showed that MR-Robin can achieve reasonable performance with only a small number of IVs (as small as 3) when the effects of invalid IVs are weak to moderate. MR-Robin allows for correlated IVs and accounts for LD among SNPs. Given most genes in the genome have only limited numbers of eQTLs, MR-Robin is particularly useful in studying the causal effects of gene expression on various complex traits.

There are some caveats of the current work and aspects that could be potentially improved in future research. First, we analyzed 13 GTEx brain tissue types as a multi-condition mediation analysis. It should be noted that the brain tissues from GTEx have only modest sample size, and a future multi-condition mediation analysis could benefit from the use of eQTL summary statistics from other studies, tissue types, and cellular conditions. Second, using CCmed_gene_, we mapped and replicated only gene-level but not SNP-level trans-associations from a cis-gene to a trans-gene in the genome. There are two main reasons: an individual SNP’s trans-effects may not be robust and replicable across conditions; and a decent proportion of SNPs may not be genotyped nor well imputed in all studies when conducting integrative analysis or replication analysis using data from several studies. Third, findings from both CCmed_gene_ and CCmed_GWAS_ should be interpreted as association rather than causation. As such, to further conduct fine-mapping of causal mediators (i.e., cisgene expression levels modulating the trans-expression or complex trait variation), additional analysis are needed. For this purpose, we proposed MR-Robin as a fine-mapping and validation method that can be used to assess the causal effect of a suspected gene on complex trait. It should be noted that even though MR-Robin can be applied to genes with as small as three eQTLs as correlated IVs, still a large proportion of the genes in the genome may have fewer than three eQTLs and are not applicable. In future work, MR-Robin can be improved in at least two aspects: the selection criteria of IVs can be further explored; and by using robust regression rather than generalized linear regression, MR-Robin may allow more inclusion of invalid IVs with very strong effects.

The R packages CCmed and MrRobin are available at https://github.com/kjgleason/.

## Methods

### Obtaining summary statistics of gene-level cis-association and conditional correlation in each single tissue type for CCmed_gene_

To detect the trios (***L**_i_*, *C_i_*, *T_j_*)’s with non-zero trans-association and mediation effects ***L**_i_* → *C_i_* → *T_j_*, two conditions both need to be satisfied – a cis-association between the eQTL SNP set ***L**_i_* and cis-gene expression *C_i_*; and a conditional correlation of cis- and trans-expression levels (*C_i_* and *T_j_*), conditioning on the eQTL genotypes ***L**_i_*. Note that here ***L**_i_* is a set of pre-selected cis-eQTLs.

In a single tissue type *k* (*k* = 1, …, *K*), a test of cis-association between ***L**_i_* and *C_i_* is conducted based on the following regression:

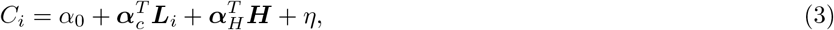

where ***α**_c_* are the cis-association effects of interest, ***H*** are covariates, ***α**_H_* are coefficients of covariates, and *η* is the error term. In the current analyses, ***H*** includes gender, genotype Principal Components (PCs), PEER factors [36] capturing expression heterogeneity, and genotyping platform. We obtain the *F*-statistics, *F_ik_*, for testing gene-level cis-association of gene *i* in tissue *k*, *H*_0_ : ***α**_c_* = **0** vs. *H_A_* : ***α**_c_* ≠ **0**.

To test for non-zero conditional correlation effects between cis- and trans-expression levels conditioning on eQTL genotypes, we perform the following regression in each single tissue type:

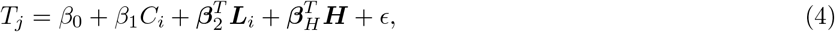

where *β*_1_ captures the conditional correlation effect of interest. We obtain the *t*-statistic, *Z_ijk_*, for cis-gene *i* and trans-gene *j* in testing *H*_0_ : *β*_1_ = 0 versus *H_A_* : *β*_1_ ≠ 0 in tissue type *k*. Note that we allow for potential partial mediation, i.e. horizontal pleiotropic effect, depicted by the direct effect ***β***_2_ in equation (4).

In the following cross-tissue (cross-condition) mediation analysis, the inputs are the two sets of the summary statistics: (i) {*F_ik_*} for testing cis-associations of all cis-genes (*i* = 1, …, *M*) in each of the *K* tissue types, and (ii) {*Z_ijk_*} for conditional correlations between *C_i_* and *T_j_* conditional on ***L**_i_* in each of the *K* tissue types.

### CCmed_gene_ for detecting cross-tissue gene-level cis-mediated trans-associations

For each trio (***L**_i_*, *C_i_*, *T_j_*), we calculate the probability that *C_i_* mediates the effects of ***L**_i_* on *T_j_* in at least *K*_1_ tissue types, P_med,*ij*_, as follows:

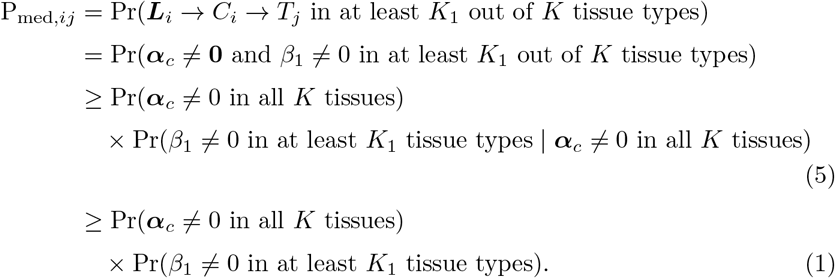

The first probability in (5) describes the probability of a gene *i* having at least one cis-eQTL in all *K* tissue types, and the lead cis-eQTL may differ by tissue type. Given robust cross-tissue cis-associations, the second probability quantifies the probability of non-zero cis-trans gene expression correlations conditioning on eQTL genotypes in at least *K*_1_ tissue types. The derivation from (5) to (1) is based on the assumption that genes with cis-associations across all tissues are more likely to affect downstream genes [25]. The probability product in (1) provides a lower (i.e., conservative) bound estimate for the probability of mediation, P_med,*ij*_.

To estimate Pr(***α**_c_* ≠ 0 in all *K* tissues), we apply a recently developed integrative association analysis approach, Primo [15, 21]. Primo takes as input the *M* × *K* matrix of cis-association summary statistics {*F_ik_*}, considers all 2^*K*^ possible cross-tissue association patterns and estimates the density function for each pattern, and returns the probabilities of 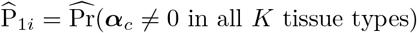 for gene *i* = 1, …, *M*. For each cis-gene *i*, to estimate the probabilities of conditional correlations for the cis-genes *i* and its 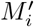 trans-genes in at least *K*_1_ tissue types, we apply the Primo algorithm to the 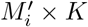 matrix of conditional correlation statistics 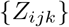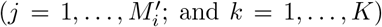, and obtain the probabilities 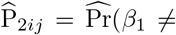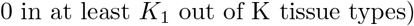. For each trio (***L**_i_*, *C_i_*, *T_j_*), the cross-tissue mediation probability can be estimated as 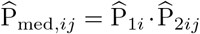. Algorithm 1 provides a summary of the algorithm.

**Algorithm 1.**
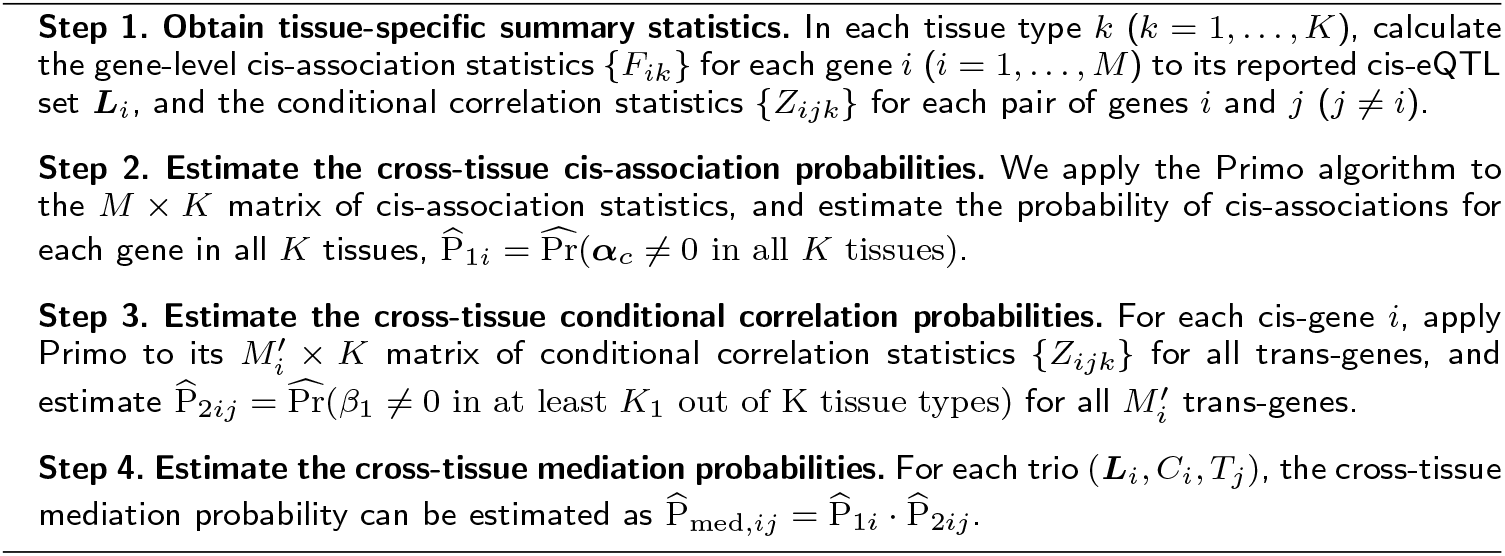
CCmed_gene_ for detecting cross-tissue gene-level cis-mediated transassociations

### CCmed_GWAS_ for detecting trans-genes associated with GWAS SNPs mediated by cis-expression levels in certain tissue-types

Different than Algorithm 1 for detecting gene-level mediation and trans-association effects, here we are interested in detecting trans-genes associated with a GWAS SNP for a complex trait or disease. It has been reported in the literature that a proportion of GWAS SNPs are also an eQTL with cis-association effects on their local gene expression levels [14, 37]. And such cis-association effects are likely to be present in some disease/trait-relevant tissue types but not all tissues. Therefore, in the mediation and trans-association analysis of GWAS SNPs, we made some tailored developments and propose the CCmed_GWAS_ algorithm.

First, we obtain the tissue-specific summary statistics for cis-association (GWAS SNPs’ eQTL associations) and conditional correlation of cis-trans gene expression.

To assess whether a GWAS SNP, *G_i_*, is associated with the cis-gene expression (*i* = 1 …, *N_G_* with *N_G_* total GWAS SNPs), we perform the following regression:

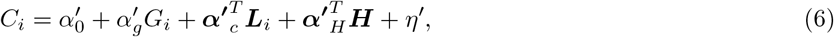

where 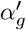 is the cis-association effect of a GWAS SNP *i* with genotype *G_i_* on cis-gene expression *C_i_*. We obtain the *t*-statistic for testing the single parameter 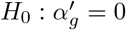 as the cis-association statistic adjusting for other GTEx-reported eQTLs. Note that here although we are interested in only the association of GWAS SNP *G_i_* to cis-gene *C_i_*, we adjust GTEx-reported eQTLs for gene *i* in the regression to reduce false associations due to LD. Also note that if an eQTL is a GWAS SNP or is in high LD (*r*^2^ > 0.5) with it, we exclude it from ***L**_i_* in the above analysis. We obtain the cis-association statistics for all *N_G_* pairs of GWAS SNPs and their cis-genes in *K* tissue types, 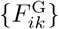. One GWAS SNP may be associated with multiple cis-genes, and those are considered as separate pairs.

For each trio of a GWAS SNP, a cis-gene and a trans-gene (*G_i_*, *C_i_*, *T_j_*), the conditional correlation statistic for the cis-gene of GWAS SNP *i* with trans-gene *j* given genotype *G_i_* in each tissue *k* can be calculated based on regression (7). Slightly different than (4), in regression (7) both eQTLs and GWAS SNP genotypes are adjusted. To test for non-zero conditional correlation effects, we perform the following regression:

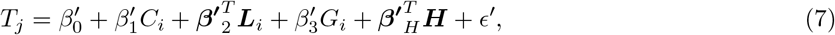

where 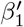 is the parameter of interest that captures the conditional correlation effect between cis and trans-expression levels, adjusting for eQTLs, the GWAS SNP and other covariates. We obtain the *t*-statistics for testing 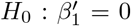 in all *K* tissue types for all trios as the conditional association statistics, 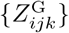.

To detect trans-genes associated with GWAS SNPs mediated by cis-genes, we propose to integrate the summary statistics from all *K* tissues to calculate the probability of mediation in at least 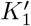 tissue types, 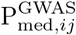. We used 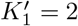 = 2 in the scz analysis.

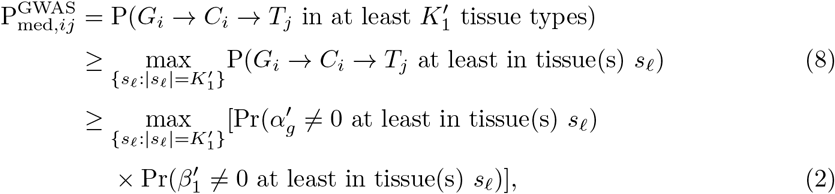

where *s*_*ℓ*_ is a set of tissue indices and is a subset of {1, 2, · · ·, *K*} with 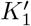 distinct tissue types. There are a total of 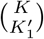 unique *s*_*ℓ*_’s. The parameters 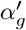 and 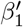 are from equations (6) and (7), with the corresponding tissue-specific test statistics 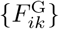 and 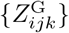 respectively.

In the above derivation, the inequality (8) follows from the fact that the probability of mediation in at least 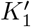 tissue types is lower bounded by the maximum probability of mediation in at least any specific set of 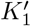 tissues. The inequality (2) holds under the assumption that cis-genes affected by GWAS SNPs are at least equally likely to affect downstream trans-genes compared to those not affected by GWAS SNPs. The maximum value of the probability products across all possible combinations of 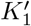 tissue types in (2) provides a lower bound estimate for the probability of a GWAS SNP *i* being associated with a trans-gene *j* via cis-mediation in at least 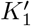 tissue types, 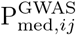. All the probabilities involved can be estimated by applying the Primo algorithm separately to the matrix of cis-association statistics and to the conditional correlation statistics matrix for each GWAS SNP *i*. The algorithm of CCmed for GWAS SNPs is summarized in Algorithm 2.

**Algorithm 2.**
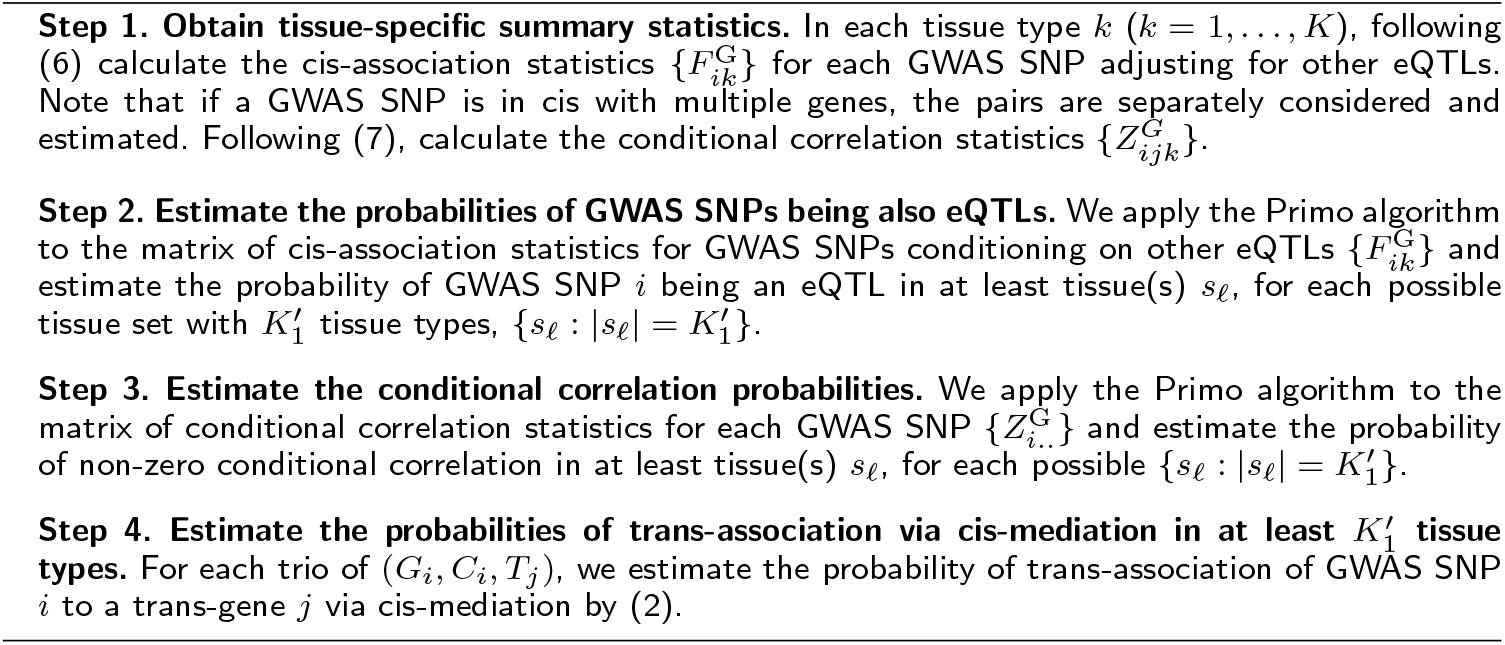
CCmed_GWAS_ for trans-genes associated with GWAS SNPs via mediation of cis-expression levels

### MR-Robin for assessing the causal effect of suspected trans-gene expression on complex trait

Many existing methods have been proposed in the Mendelian Randomization literature to assess the causal effects of a gene’s expression levels on a complex trait by harnessing Mendelian randomized genetic variants as IVs. [22, 23, 24, 25]. Earlier works assumed that genetic variants are completely mediated through gene expression [26], i.e., no “horizontal pleiotropy” [22], and are independent of unmeasured confounders of the mediator (gene expression) and the outcome (complex trait), i.e., no “correlated pleiotropy” [27]. Violations of either assumption would invalidate the SNPs as IVs and can lead to biased causal effect estimation and inference. However, it has been shown that horizontal pleiotropy is prevalent in the genome and many of the genetic effects on omics and complex traits are only partially mediated by gene expression [7, 28]. Some methods relaxed the assumptions and consider multiple SNPs as multiple IVs, while allowing some IVs to be invalid [31, 38, 30]. Those methods either require the IVs to be nearly independent or require the individual-level data, limiting the applicability in real applications. More recently, the method CAUSE [27] is proposed to account for both horizontal and correlated pleiotropy, though the method requires a large number of IVs. In contrast, many of the suspected trans-genes in our work may not have a large number of eQTLs. Another issue that is not sufficiently addressed by existing methods is the use of two samples – eQTL reference and GWAS. It is known that eQTL effects depend on cellular conditions and may not always be consistent across two samples.

To assess the causal effect of expression levels of a gene on a complex trait of interest, we propose a two-sample MR method – MR-Robin. Here we first introduce the notation. Let *β_xi_*(*i* = 1, …, *I*) denote the marginal eQTL effect of a local eQTL/IV *i* for a gene and *β_yi_* denote the marginal GWAS association effect of the eQTL/IV *i*, both from a GWAS study (though eQTL data for a GWAS study may not be available). Our goal is to test whether the effect of gene expression on the trait (γ in Figure 5) is zero, *H*_0_ : γ = 0 vs. *H_A_* : γ ≠ 0.

**Figure 5.**
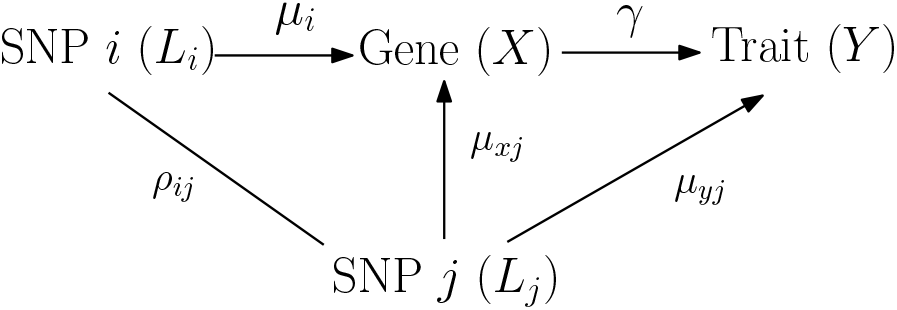
An illustration of horizontal pleiotropy in a gene region. There is a SNP *i* of interest being a candidate IV. A SNP *j* in LD with it is also an eQTL of the targeted gene and has a direct effect on the trait. When conducting MR using only marginal summary statistics, the effect of SNP *j* is not accounted for, and will confound the relationships among the SNP *i*, the gene expression and the trait. That is, horizontal (and/or correlated) pleiotropy in a gene region will bias the effect estimate based on marginal statistics for SNP *i*, without conditioning on SNP *j*.

In the presence of horizontal or correlated pleiotropy, an eQTL would be an invalid IV. And in such a case, the effect from gene to trait (γ) is not separable/identifiable from the direct effect of the eQTL nor confounding effects (for example, *μ_yj_* in Figure 5) when only the total effect estimate (marginal summary statistic) is available. The presence of horizontal or correlated pleiotropy makes it challenging to infer the effect of a gene on a trait using single-IV-based MR approaches. When there are multiple eQTLs in the gene region, as shown in Figure 5, the presence of one eQTL with horizontal or correlated pleiotropic effect would also render all eQTLs invalid if they are in LD. Specifically, consider a SNP *i* of interest with genotype *L_i_*, eQTL effect *μ_i_* and gene on trait effect of interest γ in Figure 5. A SNP *j* in LD (*ρ_ij_* ≠ 0) affects the complex trait *Y* via an independent pathway (with horizontal pleiotropic effect *μ_y_j__* ≠ 0). Since SNPs *i* and *j* are in LD, SNP *i* would also be associated with *Y* via an independent pathway (*ρ_ij_* · *μ_y_j__* ≠ 0) not mediated through gene expression *X*. Thus, if SNP *j* has horizontal pleiotropic effect, then SNP *i* in LD with it would have biased effect estimate of γ based on marginal statistics *β_yi_/β_xi_*. In the Supplemental Materials, we derive the biases of the effect estimate *β_yi_/β_xi_* for SNP *i* with respect to the true effect γ, and show that the magnitude of the bias depends on LD, eQTL and pleiotropic effect sizes, and other factors, and thus are SNP-specific. It should be noted that although the derivation is based on two SNPs *i* and *j*, the conclusion extends to multiple SNPs in a gene region: if there is horizontal or correlated pleiotropic effect for at least one SNP in a gene region, the effect estimates based on marginal summary statistics for all SNPs in LD would be biased to varying extents.

Given the bias derived for *β_yi_/β_xi_* w.r.t γ, we model that *β_yi_/β_xi_* = (γ+γ_*i*_), where γ_*i*_ denotes the SNP-specific bias. The bias is zero if there is neither horizontal nor correlated pleiotropic effect in the region. The bias is small to negligible for some eQTLs if those eQTLs themselves are valid IVs when adjusting for invalid IV *L_j_* and those eQTLs being in moderate-to-weak LD with the invalid IV(s) and when the pleiotropic effect of SNP *j* is not strong (i.e., small *ρ_ij_* · *μ_yj_*). It follows that

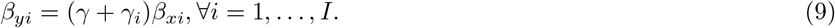

And equivalently,

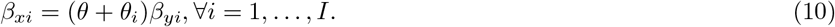

where *θ* captures the dependence between *β_xi_* and *β_yi_*, and *θ_i_* is the SNP-level deviation from the common effect *θ* in the presence of pleiotropy.

In the above equation, *β_xi_* is the marginal eQTL effect of SNP *i* to gene expression in the GWAS study and is often not available, since most GWAS studies do not have gene expression data measured. The availability of multitissue eQTL summary statistics from trait-relevant tissue types in a reference eQTL study such as GTEx provides a valuable resource to estimate *β_xi_*, given cis-eQTL effects are often replicable across studies. We model SNP *i*’s eQTL effect in tissue *k* (*k* = 1, …, *K*) in the reference multitissue eQTL data as a function of the eQTL effect in the GWAS data (*β_xi_*) and an error term. Based on (10), we propose the following model of MR-Robin for testing trait-association of a trans-gene using only summary statistics from GWAS and multitissue eQTL reference:

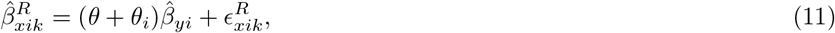

where ^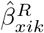^ is the eQTL effect estimate of SNP *i* in tissue *k* in the reference eQTL data, and 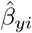 is the GWAS effect estimate for SNP *i*; and *θ* captures the shared correlation of GWAS and eQTL statistics among all SNPs and is non-zero and bounded if and only if the true mediation effect γ is non-zero and bounded; *θ_i_* represents the SNP-specific bias due to horizontal or correlated pleiotropy in the region and is a SNP-specific random-slope; and 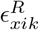 is a random error that follows a multivariate normal distribution 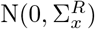. The diagonal elements of 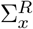 are the variance estimates of eQTL effects, 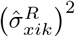’s, and the off-diagonal elements are 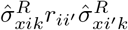 where 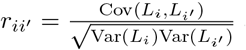 and **R** = {*r_ii′_*} is the genotype correlation matrix. In the reverse regression (11), the eQTL effect estimates from multiple tissue types, 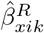, are considered as the response variable while the GWAS association effects 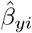 are considered as the predictor. This is mainly to take advantage of the rich information in multitissue eQTL datasets (i.e., variation in response). If there are multiple sets of correlated or independent GWAS summary statistics from the same population/ethnicity, one may not need to use reverse regression and can instead treat GWAS association effects as the response. Each observation in the regression (11) is a tissue-specific eQTL effect, with a total of *I* × *K* (sample-by-tissue) observations. By testing the shared correlation of tissue-specific eQTL and the corresponding GWAS association effects for all eQTLs in the same gene (*H*_0_ : *θ* = 0 vs. *H_A_* : *θ* ≠ 0) while also allowing for SNP-level deviation, we can test the effect of gene expression on trait (*H*_0_ : γ = 0 vs. *H_A_* : γ ≠ 0), allowing invalid and correlated IVs.

Note that many existing methods in the MR literature [27, 30, 38] include an intercept or a random intercept in the model to capture the direct effect from genotype to trait, i.e. horizontal pleiotropy. In contrast, in the MR-Robin model, there is no intercept nor random intercept. Instead, we include a random slope for each SNP to capture the effect due to potential pleiotropy in the region. This is because, by allowing correlated IVs and considering all eQTLs in a region, as shown above when there is a non-zero pleiotropic effect, most of the SNPs in the LD region would be affected with a non-zero (but possibly negligible) SNP-specific deviation *θ_i_*. Allowing correlated IVs and some invalid IVs even when the number of IVs are limited is also a major innovation of our model. Due to limited numbers of eQTLs/IVs for most genes in the genome, a model with both an intercept and a random slope may not be identifiable and thus are not explored.

To account for uncertainty in the eQTL effect estimation, we perform a weighted mixed-effects regression analysis and weigh each “observation” (i.e., a tissue-specific eQ L e ect) by the reciprocal of the estimated standard error for 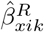, i.e., 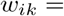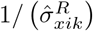. We obtain the *t*-statistic for testing the fixed effect of interest *θ* as our test statistic. To obtain the *P*-value while accounting for LD and the uncertainty in the estimation of *β_yi_*’s, we adopt a resampling based approach to generate the null test statistics. In each resampling *b* (*b* = 1, …, *B*), we sample a vector of GWAS effects from a multivariate distribution, 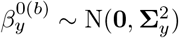, where the diagonal and off-diagonal elements are 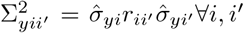 with *r_ii′_* being the genotype correlation matrix and 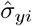 being the estimated standard error for 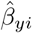. We apply the same weighted model (11) on data 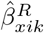’s and 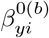’s to obtain a null statistic. We repeat the resampling process at least *B* = 10, 000 times and calculate the *P*-value. The MR-Robin algorithm is summarized in Algorithm 3.

**Algorithm 3.**
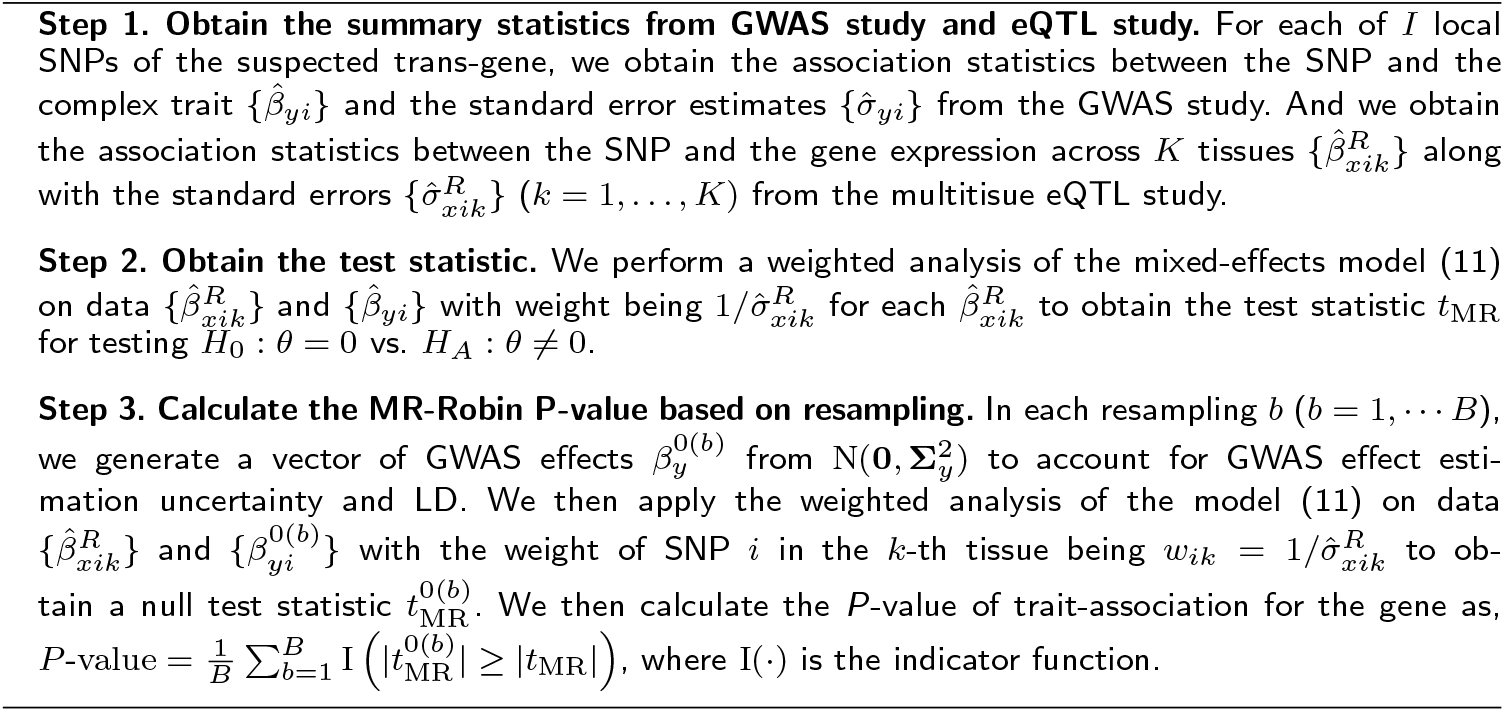
MR-Robin for assessing the causal effect of gene expression of a gene on complex trait with summary statistics from GWAS and multitissue eQTL study

## Abbreviations

GWAS: genome-wide association studies
SNP: single nucleotide polymorphism
QTL: quantitative trait locus
eQTL: expression quantitative trait locus
Primo: Package in R for Integrative Multi-Omics association analysis
LD: linkage disequilibrium
GTEx: Genome-Tissue Expression project
CMC: CommonMind Consortium
MR: Mendelian randomization
IV: Instrumental variables
PGC: Psychiatric Genomics Consortium
scz: Schizophrenia

## Acknowledgements

We thank the GTEx Consortium. We thank Drs. Francois Aguet, Kristin Ardlie for providing cis-eQTL summary statistics.

## Availability of data and material

The R package CCmed is available at https://github.com/kjgleason/CCmed. The R package MrRobin is available at https://github.com/kjgleason/MrRobin.

## Author’s contributions

LSC and FY conceived the project and developed the methods. LSC, FY and KJG wrote the manuscript. LSC and KJG analyzed the data. JW, JD, XH and BLP provided valuable suggestions to the development of the methods and analysis. KJG and FY developed the R software package.

## Ethics approval and consent to participate

Not applicable.

## Consent for publication

Not applicable.

