## Supplemental Materials for "CCmed: cross-condition mediation analysis for identifying robust trans-eQTLs and assessing their effects on human traits"

METHOD

#### Supplemental Methods

Bias in $\beta_{yi}/\beta_{xi}$ as an estimand for $\gamma$ in the presence of invalid instrumental variables (IVs) being in LD

Here we present details on the derivation of the bias in the ratio of marginal GWAS association effect to marginal eQTL effect for a SNP $i$ as an estimand for the effect of the trans-gene on the trait, $\gamma$, in the presence of SNP(s) in LD and with horizontal pleiotropy effects. We will show that the bias is SNP-specific. Without loss of generality, we assume that there are two SNPs $i$ and $j$ in LD, and SNP $i$ is a valid IV if conditioning on SNP $j$ with a horizontal pleiotropic effect as depicted in Figure 5 of the main text. For multiple eQTLs in an LD block, one can consider them as being conditionally valid IVs and invalid IVs. Below are the data generating models in a GWAS study:

$$X = \mu_{x0} + \mu_i L_i + \mu_{xj} L_j + \epsilon_x, \tag{S1}$$

$$Y = \mu_{y0} + \gamma X + \mu_{yj} L_j + \epsilon_y, \tag{S2}$$

where $X$ is the gene expression levels and $Y$ is the continuous complex trait of interest in a GWAS study; and $L_i$ and $L_j$ are the genotypes for SNPs $i$ and $j$, respectively. As a valid IV given $L_j$, the genotype of SNP $i$ ($L_i$) is independent of the error terms $\epsilon_x$ and $\epsilon_y$. In the above models, the conditional association between $X$ and $L_i$ given $L_j$ is captured by $\mu_i$, and the conditional association between $Y$ and $L_i$ given $L_j$ is $\gamma \cdot \mu_i$. And the ratio of the two, $\frac{\gamma\mu_i}{\mu_i}$, recovers the true effect of interest, $\gamma$.

Without adjusting for SNP $j$, the summary statistics are calculated based on the following marginal models:

$$X = \beta_{x0} + \beta_{xi} L_i + \epsilon'_x, \tag{S3}$$

$$Y = \beta_{y0} + \beta_{yi} L_i + \epsilon'_y, \tag{S4}$$

where $\beta_{xi}$ and $\beta_{yi}$ are the marginal eQTL and GWAS association effects, respectively, in the GWAS study. Note that one could also adjust covariates in the above models (S1)-(S4) and that does not affect our conclusion. We ignore covariates for simplicity. Define $\rho_{ij} = \frac{\text{Cov}(L_i,L_j)}{\text{Var}(L_i)}$, in terms of parameters in (S1) and (S2), the marginal effects $\beta_{xi} = \frac{\text{Cov}(X,L_i)}{\text{Var}(L_i)} = \frac{\text{Cov}(\mu_{x0}+\mu_i L_i+\mu_{xj}L_j+\epsilon_x,L_i)}{\text{Var}(L_i)} = \mu_i + \mu_{xj}\rho_{ij}$, and $\beta_{yi} = \frac{\text{Cov}(Y,L_i)}{\text{Var}(L_i)} = \frac{\text{Cov}(\mu_{y0}+\gamma X+\mu_{yj}L_j+\epsilon_y,L_i)}{\text{Var}(L_i)} = [\gamma + (\gamma\mu_{xj} + \mu_{yj})\frac{\rho_{ij}}{\mu_i}]\mu_i$.

It can be seen that the bias of marginal eQTL effect estimate for SNP $i$ on gene expression, $\beta_{xi}$, with respect to the true eQTL effect, $\mu_i$, is $\mu_{xj}\rho_{ij}$. And the bias of marginal GWAS effect estimate for SNP $i$ on complex trait, $\beta_{yi}$, with respect to the mediated effect from SNP to gene to trait, $\gamma\mu_i$, is $(\gamma\mu_{xj} + \mu_{yj})\rho_{ij}$. And it can be derived that the bias of the ratio of marginal GWAS to eQTL effect estimates, $\beta_{yi}/\beta_{xi}$, with respect to the true effect, $\gamma$, is given by $\frac{\mu_{yj}\rho_{ij}}{\mu_i+\mu_{xj}\rho_{ij}}$. All the biases are functions of SNP $i$'s eQTL effect size, LD strength to SNP $j$ and effect size of the pleiotropy. Therefore, the bias will vary from SNP to SNP.

#### Supplemental Results

##### Simulation studies to evaluate the performance of CCmed

Here we report additional details regarding the simulation studies evaluating the performance of the CCmed algorithm.

*The performance of CCmed$_{gene}$ in identifying robust gene-level trans-associations*

Here we describe additional details of the simulation evaluating the performance of CCmed$_{\text{gene}}$ in identifying robust gene-level trans-associations (results of the simulations are presented in Table 1A in the main text). In this simulation, we generated 5000 genotypes for SNPs and grouped each of 10 SNPs as the cis-eQTL set for a cis-gene to generate 500 cis-eQTL sets for 500 cis-genes. In each cis-eQTL set, the pairwise correlations between SNPs were set to be 0.3. Based on the genotypes, in each tissue type, we randomly selected 1 SNP as the causal eSNP to generate the cis- and trans-gene expression levels. Note that this way, the causal eSNPs varied across tissues. For each pair of a cis-eQTL set and cis-gene, we generated 500 trans-gene expression levels. We generated cis-trans gene expression data from 10 correlated tissue types. The proportions of trios with the SNP set being associated with cis-gene in all 10 tissue types, in each combination of exactly 9 tissue types (there are 10 of them), and in each combination of exactly 8 tissue types (there are 45 of them) were set to be 0.124, 0.026, and 0.008, respectively. The proportion of trios with the cis-eQTL set being associated with cis-gene in none of the tissues was 0.216, and the probabilities for each of the rest of the possible association patterns were set to be the same. Among the trios with cis-associations in all 10 tissues, 60% of them were simulated with conditional cis-trans gene expression correlations in at least 9 tissues. Among trios with cis-associations in exactly 9 or 8 tissue types, 60% of them had non-zero conditional cis-trans gene expression correlations in exactly the same tissue types as their corresponding cis-association tissue types. And in the simulation studies, we are interested in detecting the trios with cis-association and conditional expression correlation in at least 9 out of 10 tissue types. For the rest of the trios, the conditional cis-trans association patterns were randomly generated with a probability of associations in none of the tissues to be 0.4885, and with probabilities in each of the rest of the possible patterns being 0.0005. Among those trios with non-zero cis-mediated trans-associations, 50% of them also had a non-zero direct effect from SNPs on the trans-gene expression levels. Nonzero cis-association and conditional cis-trans association effect sizes were generated from multivariate normal distribution with means of either a vector of 0.8 or −0.8, standard deviations 0.3 and correlations 0.3 across tissues. The effect sizes for direct effects were generated from a normal distribution with mean 0 and standard deviation 0.3. This simulation setup mimics weak total trans-associations (note that the mean of each nonzero total trans-association is of size $0.8 \times 0.8$) observed in the GTEx study. Performance of $\text{CCmed}_{\text{gene}}$ in detecting gene-level trans-associations mediated by cis-gene expression in at least $K_1 = 9$ out of the 10 tissue types is presented in Table 1A in the main text.

*The performance of $CCmed_{GWAS}$ in identifying cis-mediated trans-genes for one (GWAS) SNP in selected tissue-types*

Here we describe additional details of the simulation evaluating the performance of $\text{CCmed}_{\text{GWAS}}$ in identifying cis-mediated trans-genes for a GWAS SNP in selected tissue-types (results of the simulations are presented in Table 1B in the main text). In this simulation, we simulated cis-gene expression levels being affected by 3 correlated eQTLs with correlation 0.3. We focused on one of them as the (GWAS) SNP of interest and generated the trans-gene expression levels being affected by the SNP in selected tissue types. The proportion of trios with the SNP being associated with cis-gene expression in none of the tissue types, in each combination of exactly 1 tissue (there are 10 of them), exactly 2 tissues (there are 45 of them), and exactly 3 tissues (there are 120 of them) were 0.298, 0.01, 0.006, and 0.002, respectively. And the proportions for each of the rest of the possible association patterns were all the same. Among the trios with cis-association in exactly 1 tissue type, exactly 2 tissue types and exactly 3 tissue types, the proportions of them that had non-zero conditional cis-trans expression correlations in the same tissue types were 60%. For the rest of the trios, the conditional cis-trans expression correlations were randomly generated with a probability of non-zero correlations in none of the tissues to be 0.4885, and with probabilities in each of the rest of the possible patterns being 0.0005. Same as in the previous simulation, among those trios with non-zero cis-mediated trans-associations, 50% of them also had a non-zero direct effect of the SNP on the trans-gene. Nonzero cis-association and conditional cis-trans expression correlation effect sizes were generated from multivariate normal distributions with means of either a vector of 1 or −1, standard deviations 0.5 and correlations 0.3 across tissues. The effect sizes for direct effects were generated from a normal distribution with mean 0 and standard deviation 0.5. This simulation considers scenarios with weak to moderate effects in certain tissue types. Performance of $\text{CCmed}_{\text{GWAS}}$ in identifying associations between the GWAS SNP and trans-gene mediated by cis-gene expression in at least $K_1' = 2$ tissues is presented in Table 1B in the main text.

#### Simulation studies to evaluate the performance of MR-Robin

*Data generation*

We evaluated the performance of MR-Robin using simulations. In each simulation scenario, we simulated data for a total of $N = N_g + N_R = 10,300$ independent subjects: $N_g = 10,000$ subjects in a GWAS study, and $N_R = 300$ subjects in a reference multitissue eQTL study of $K = 10$ tissues.

First, we simulated an $N \times I$ genotype matrix for each gene, $\mathbf{L}$, comprised of $Q$ independent LD blocks with 20 SNPs in each block (thus, a total of $I = 20 \times Q$ SNPs for each gene). From each LD block, we selected 1 SNP to be the true eQTL. The $N_g \times Q$ genotype matrix of the $Q$ true eSNPs in the GWAS study is denoted $\mathbf{G}$, and we generated phenotypes in the GWAS study according to the following data generation models:

$$X = \mathbf{G}\boldsymbol{\mu_x} + \epsilon_x, \tag{S5}$$

$$Y = \gamma X + \sum_{q=1}^{Q} \mu_{yq}\mathbf{g}_q + \epsilon_y, \tag{S6}$$

In model S5, $X$ is a vector of gene expression levels; $\mathbf{G}$ are the genotypes of eSNPs; $\boldsymbol{\mu_x} \sim N_Q(\mathbf{0}, 0.25 \cdot \mathbf{I})$ are the eQTL effects of eSNPs from independent LD blocks; and $\epsilon_x \sim N(0, 0.25)$ are error terms. In model S6, $Y$ is a vector of the complex trait; $\gamma$ is the parameter of interest – the effect of gene $X$ on trait $Y$ – with $\gamma = 0$ under the null and $\gamma = 0.25$ under the alternative; $\mathbf{g}_q$ is the genotype vector of SNP $q$; $\mu_{yq}$ is the direct effect of SNP $q$ on $Y$; and $\epsilon_y \sim N(0, 1)$ are the error terms. When SNP $q$ is a valid IV, the direct effect on Y is $\mu_{yq} = 0$; otherwise, $\mu_{yq} \sim N(0, 0.05)$. Across scenarios we vary the proportion of the $Q$ SNPs that are invalid.

Data from the eQTL study was generated based on the model:

$$\mathbf{X}^{\mathrm{R}} = \mathbf{G}^{\mathrm{R}}\boldsymbol{\mu}_{\boldsymbol{x}}^{\mathrm{R}} + \epsilon_x^{\mathrm{R}}, \tag{S7}$$

where $\mathbf{X}^{\mathrm{R}}$ is an $N_R \times K$ matrix of expression levels measured in $K$ tissues; $\mathbf{G}^{\mathrm{R}}$ is a $N_R \times Q$ genotype matrix of $Q$ eSNPs in the eQTL study; $\boldsymbol{\mu}_{\boldsymbol{x}}^{\mathrm{R}}$ is a $Q \times K$ matrix of the tissue-specific eQTL effects; and $\epsilon_x^{\mathrm{R}} \sim N(0, 1)$ are the error terms. Each column of $\boldsymbol{\mu}_{\boldsymbol{x}}^{\mathrm{R}}$ is independently drawn from $N_Q(\boldsymbol{\mu_x}, 0.05 \cdot \mathbf{I})$, where $\boldsymbol{\mu_x}$ is from model S5.

*Summary statistics*

After individual-level data was generated in each simulation, we calculated the marginal eQTL and GWAS summary statistics, and obtained the marginal effect estimate of each SNP $i$ on gene expression in tissue $k$ in the reference eQTL study, $\beta_{xik}^{R}$; and the marginal effect estimate of each SNP $i$ on its simulated trait in the GWAS study, $\beta_{yi}$, for two-sample MR analyses. We also obtained the standard error estimates for marginal eQTL and GWAS effects.

*Description of competing two-sample MR models and methods*

Finally, we applied MR-Robin to the summary statistics $\hat{\beta}_{xik}^{R}$ and $\hat{\beta}_{yi}$ and their standard errors, and obtained the $P$-value for each simulated gene, as described in Algorithm 3 in the main text.

For comparison, we included three competing models. The first one is a single-tissue model with GWAS effects as the response and eQTL effects as the predictor. No intercept is included. Each observation is weighted by $1/\sigma_{yi}^2$. We selected one tissue at random from all simulated tissues for the model, and obtained the parametric Wald $P$-values testing the hypotheses $H_0 : \gamma = 0$ vs. $H_A : \gamma \neq 0$.

The second model extends the above single-tissue model to a multitissue model with multitissue eQTL effects as predictor and the corresponding GWAS effects as response, without an intercept:

$$\hat{\beta}_{yi} = \gamma\hat{\beta}^{R}_{xik} + \epsilon \tag{S10}$$

Each observation is weighted by $1/\sigma^2_{yi}$. We obtain the test statistics for testing the hypotheses $H_0 : \gamma = 0$ vs. $H_A : \gamma \neq 0$, and calculate the $P$-values by resampling to account for the correlation among tissues and LD.

As a third comparison model, we performed a weighted, random-intercept regression based on the following reverse-regression model:

$$\hat{\beta}^{R}_{xik} = \theta\hat{\beta}_{yi} + \mu_i + \varepsilon, \tag{S11}$$

where $\mu_i$ is the SNP-specific random intercept for each IV with mean zero. We test the hypotheses $H_0 : \theta = 0$ vs. $H_A : \theta \neq 0$. To make a fair comparison, we weighted each observation by $1/\sigma^2_{xik}$. We estimated $P$-values based on resampling.

We also compared the performance of MR-Robin to three existing Mendelian randomization methods reported in the literature: MR-RAPS [1], MR-Egger [2], and MRMix [3]. Note that these methods were developed for settings where many, independent genetic variants may be available as candidate IVs, and those methods are all developed for single-tissue eQTL statistics. Therefore, they may not be expected to perform well in the currently proposed setting, where a limited number of correlated variants are available as candidate IVs (i.e. variants in cis with a particular gene). Nonetheless, we include the methods for comparison. For each method, we performed the analysis using eQTL statistics from a single tissue type – the same tissue type selected for the single-tissue model (the first competing model described above).

*MR-Robin controls type I error rate with moderate proportion of invalid IVs*

In Scenario 1, we evaluated the robustness of MR-Robin to the proportion of invalid IVs. We simulated the data using $Q = 10$ LD blocks, varying the proportion of invalid IVs across settings. That is, we varied the proportion of eSNPs having direct effects on the complex trait $Y$ (i.e. effects not mediated through gene expression $X$). Over 10,000 simulations, we compare the type I error rate and power of MR-Robin to competing methods. $P < 0.05$ was used as the significance criterion for each method. Tables S1, S2 and S3 compare the methods when the selection LD $r^2$ threshold is set to 0.8, 0.3, and 0.01, respectively (results using selection LD $r^2$ threshold of 0.5 are reported in Table 1 in main text).

Based on the results, we observe that competing methods are generally unable to control the type I error rate when there are any invalid IVs and IVs are in LD. On the other hand, MR-Robin is able to control the type I error rate when a majority of IVs are valid (e.g. when up to 30% are invalid). Power is reasonable for all methods when a majority of IVs are valid.

Since our method allows for correlated IVs and it is hard to define invalid versus valid IVs when SNPs are correlated, the proportions of valid IVs in the tables are the proportion of LD blocks with no pleiotropy, and is only an approximation of the valid IVs among all selected ones. In each table, we also presented the average numbers of selected IVs that are from valid versus invalid LD blocks.

**Table S1 Simulation results evaluating the performance of MR-Robin. Averaged type I error rates and power over 10,000 simulations are shown by percentage of valid instruments. 10 LD blocks were simulated, with one true eQTL per LD block. Instruments were selected sequentially: the eSNP with the strongest association with gene expression was selected, and the next selected eSNP is the strongest-associated SNP remaining also with LD $r^2 < 0.8$ with any already-selected eSNPs.**

| Method | Proportion of Valid IVs (%) 100 | 90 | 80 | 70 | 60 | 50 | 30 |
|---|---|---|---|---|---|---|---|
| | Type I error rate | | | | | | |
| MR-Robin | 0.050 | 0.064 | 0.072 | 0.091 | 0.114 | 0.140 | 0.213 |
| A single tissue MR model with no intercept | 0.466 | 0.509 | 0.532 | 0.554 | 0.574 | 0.590 | 0.588 |
| A multitissue MR model with a fixed slope and no intercept | 0.048 | 0.075 | 0.093 | 0.111 | 0.129 | 0.140 | 0.150 |
| Random Intercept | 0.048 | 0.074 | 0.092 | 0.113 | 0.130 | 0.140 | 0.150 |
| MR-RAPS | 0.431 | 0.701 | 0.835 | 0.899 | 0.927 | 0.937 | 0.940 |
| MR-Egger | 0.257 | 0.332 | 0.379 | 0.421 | 0.440 | 0.454 | 0.472 |
| MRMix | 0.164 | 0.221 | 0.275 | 0.322 | 0.381 | 0.425 | 0.515 |
| | Power | | | | | | |
| MR-Robin | 0.985 | 0.943 | 0.902 | 0.854 | 0.803 | 0.760 | 0.647 |
| A single tissue MR model with no intercept | 0.996 | 0.979 | 0.960 | 0.940 | 0.917 | 0.900 | 0.864 |
| A multitissue MR model with a fixed slope and no intercept | 0.999 | 0.948 | 0.888 | 0.824 | 0.773 | 0.718 | 0.618 |
| Random Intercept | 0.999 | 0.950 | 0.890 | 0.828 | 0.780 | 0.724 | 0.618 |
| MR-RAPS | 1.000 | 0.997 | 0.994 | 0.991 | 0.986 | 0.981 | 0.974 |
| MR-Egger | 0.912 | 0.856 | 0.796 | 0.750 | 0.715 | 0.696 | 0.640 |
| MRMix | 0.537 | 0.527 | 0.537 | 0.530 | 0.535 | 0.542 | 0.561 |
| | Avg number of SNPs selected (valid/invalid) | | | | | | |
| All methods | 62.1/0 | 55.8/6.1 | 49.6/12.4 | 43.3/18.6 | 36.9/25.0 | 30.8/31.1 | 18.4/43.5 |

**Table S2 Simulation results evaluating the performance of MR-Robin. Averaged type I error rates and power over 10,000 simulations are shown by percentage of valid instruments. 10 LD blocks were simulated, with one true eQTL per LD block. Instruments were selected sequentially: the eSNP with the strongest association with gene expression was selected, and the next selected eSNP is the strongest-associated SNP remaining also with LD $r^2 < 0.3$ with any already-selected eSNPs.**

| Method | Proportion of valid IV (%) 100 | 90 | 80 | 70 | 60 | 50 | 30 |
|---|---|---|---|---|---|---|---|
| | Type I error rate | | | | | | |
| MR-Robin | 0.049 | 0.055 | 0.060 | 0.067 | 0.076 | 0.080 | 0.108 |
| A single tissue MR model with no intercept | 0.122 | 0.169 | 0.194 | 0.210 | 0.224 | 0.234 | 0.244 |
| A multitissue MR model with a fixed slope and no intercept | 0.050 | 0.069 | 0.081 | 0.100 | 0.108 | 0.117 | 0.124 |
| Random Intercept | 0.051 | 0.066 | 0.076 | 0.093 | 0.101 | 0.111 | 0.117 |
| MR-RAPS | 0.118 | 0.548 | 0.749 | 0.843 | 0.882 | 0.896 | 0.878 |
| MR-Egger | 0.055 | 0.124 | 0.155 | 0.180 | 0.187 | 0.195 | 0.197 |
| MRMix | 0.177 | 0.250 | 0.323 | 0.379 | 0.419 | 0.464 | 0.530 |
| | Power | | | | | | |
| MR-Robin | 0.950 | 0.893 | 0.827 | 0.757 | 0.687 | 0.627 | 0.480 |
| A single tissue MR model with no intercept | 0.981 | 0.924 | 0.869 | 0.810 | 0.767 | 0.717 | 0.640 |
| A multitissue MR model with a fixed slope and no intercept | 0.998 | 0.941 | 0.875 | 0.805 | 0.746 | 0.688 | 0.580 |
| Random Intercept | 0.998 | 0.939 | 0.872 | 0.801 | 0.741 | 0.686 | 0.572 |
| MR-RAPS | 0.999 | 0.995 | 0.988 | 0.982 | 0.975 | 0.969 | 0.953 |
| MR-Egger | 0.821 | 0.704 | 0.625 | 0.553 | 0.506 | 0.476 | 0.398 |
| MRMix | 0.576 | 0.575 | 0.565 | 0.574 | 0.578 | 0.586 | 0.594 |
| | Avg number of SNPs selected (valid/invalid) | | | | | | |
| All methods | 16.6/0 | 14.9/1.7 | 13.2/3.3 | 11.6/5.0 | 9.9/6.7 | 8.3/8.3 | 4.9/11.6 |

*MR-Robin controls type I error rate with small number of IVs*

In Scenario 2, we evaluated the performance of MR-Robin when the number of selected IVs is small. We simulated the data using $Q = 3$ LD blocks, with two blocks without pleiotropy and one block with pleiotropy (thus the proportion of LD blocks with pleiotropic effects is fixed at 33.3%). Table S4 shows the type I error rates and power when the selection LD $r^2$ threshold is set to 0.8, 0.5, 0.3, 0.2, 0.1 and 0.01. As shown in the table, MR-Robin performs reasonably well even when the number of IVs is very limited. Though in this setting, MR-Robin requires the IVs to be less dependent ($r^2 < 0.3$). MR-Robin outperforms competing methods in this setting.

**Table S3 Simulation results evaluating the performance of MR-Robin. Averaged type I error rates and power over 10,000 simulations are shown by percentage of valid instruments. 10 LD blocks were simulated, with one true eQTL per LD block. Instruments were selected sequentially: the eSNP with the strongest association with gene expression was selected, and the next selected eSNP is the strongest-associated SNP remaining also with LD $r^2 < 0.01$ with any already-selected eSNPs.**

| Method | Proportion of valid IV (%) | | | | | | |
|---|---|---|---|---|---|---|---|
| | 100 | 90 | 80 | 70 | 60 | 50 | 30 |
| | Type I error rate | | | | | | |
| MR-Robin | 0.050 | 0.048 | 0.046 | 0.044 | 0.045 | 0.043 | 0.041 |
| A single tissue MR model with no intercept | 0.046 | 0.043 | 0.046 | 0.046 | 0.054 | 0.053 | 0.059 |
| A multitissue MR model with a fixed slope and no intercept | 0.048 | 0.039 | 0.033 | 0.040 | 0.045 | 0.046 | 0.054 |
| Random Intercept | 0.049 | 0.038 | 0.032 | 0.036 | 0.041 | 0.043 | 0.052 |
| MR-RAPS | 0.044 | 0.434 | 0.659 | 0.785 | 0.843 | 0.860 | 0.865 |
| MR-Egger | 0.037 | 0.088 | 0.109 | 0.128 | 0.132 | 0.137 | 0.139 |
| MRMix | 0.183 | 0.281 | 0.363 | 0.425 | 0.476 | 0.530 | 0.607 |
| | Power | | | | | | |
| MR-Robin | 0.920 | 0.812 | 0.714 | 0.615 | 0.518 | 0.444 | 0.290 |
| A single tissue MR model with no intercept | 0.883 | 0.751 | 0.646 | 0.561 | 0.495 | 0.442 | 0.341 |
| A multitissue MR model with a fixed slope and no intercept | 0.995 | 0.880 | 0.778 | 0.687 | 0.612 | 0.547 | 0.417 |
| Random Intercept | 0.995 | 0.878 | 0.773 | 0.681 | 0.597 | 0.539 | 0.401 |
| MR-RAPS | 0.999 | 0.991 | 0.981 | 0.976 | 0.968 | 0.960 | 0.948 |
| MR-Egger | 0.577 | 0.478 | 0.402 | 0.352 | 0.318 | 0.290 | 0.237 |
| MRMix | 0.709 | 0.700 | 0.697 | 0.686 | 0.688 | 0.699 | 0.701 |
| | Avg number of SNPs selected (valid/invalid) | | | | | | |
| All methods | 6.6/0 | 5.9/0.7 | 5.2/1.3 | 4.6/2.0 | 3.9/2.6 | 3.3/3.3 | 2.0/4.6 |

**Table S4 Simulation results evaluating the performance of MR-Robin when there is a small number of IVs. Averaged type I error rates and power over 10,000 simulations are shown by IV selection criteria. 3 LD blocks were simulated, with two blocks without pleiotropic effects (valid IVs) and one block with (invalid IV). Results shown for six IV selection criteria (LD $r^2 <$ 0.8, 0.5, 0.3, 0.2, 0.1, and 0.01).**

| Method | LD selection criteria ($r^2$) | | | | | |
|---|---|---|---|---|---|---|
| | $0.8$ | $0.5$ | $0.3$ | $0.2$ | $0.1$ | $0.01$ |
| | Type I error rate | | | | | |
| MR-Robin | 0.129 | 0.113 | 0.070 | 0.049 | 0.030 | 0.011 |
| A single tissue MR model with no intercept | 0.571 | 0.441 | 0.239 | 0.135 | 0.080 | 0.033 |
| A multitissue MR model with a fixed slope and no intercept | 0.154 | 0.144 | 0.114 | 0.067 | 0.044 | 0.012 |
| Random Intercept | 0.156 | 0.147 | 0.114 | 0.074 | 0.053 | 0.023 |
| MR-RAPS | 0.780 | 0.727 | 0.665 | 0.659 | 0.668 | 0.720 |
| MR-Egger | 0.436 | 0.330 | 0.231 | 0.208 | 0.213 | 0.259 |
| MRMix | 0.298 | 0.306 | 0.319 | 0.318 | 0.304 | 0.303 |
| | Power | | | | | |
| MR-Robin | 0.686 | 0.638 | 0.482 | 0.410 | 0.330 | 0.202 |
| A single tissue MR model with no intercept | 0.835 | 0.757 | 0.550 | 0.432 | 0.314 | 0.150 |
| A multitissue MR model with a fixed slope and no intercept | 0.611 | 0.591 | 0.493 | 0.424 | 0.342 | 0.180 |
| Random Intercept | 0.618 | 0.596 | 0.508 | 0.445 | 0.371 | 0.229 |
| MR-RAPS | 0.958 | 0.948 | 0.928 | 0.925 | 0.925 | 0.920 |
| MR-Egger | 0.695 | 0.608 | 0.483 | 0.419 | 0.392 | 0.350 |
| MRMix | 0.512 | 0.502 | 0.523 | 0.559 | 0.556 | 0.585 |
| | Avg # of SNPs selected | | | | | |
| All methods | 18.0/8.4 | 9.6/4.6 | 4.2/2.0 | 3.1/1.5 | 2.5/1.2 | 2.0/1.0 |

#### MR-Robin validated trans-genes showing evidence of association with scz

In Table S5, we present detailed information on the 46 trans-genes for scz-GWAS SNPs identified by $\text{CCmed}_{\text{GWAS}}$ at 80% probability cutoff from GTEx data and validated by MR-Robin at the $P$-value cutoff of 0.05.

**Table S5 Detailed information on the 46 trans-genes for scz-GWAS SNPs, identified by $\text{CCmed}_{\text{GWAS}}$ at 80% probability cutoff from GTEx data and validated by MR-Robin at the $P$-value cutoff of 0.05**

| Validated trans-Gene | | $\text{CCmed}_{\text{GWAS}}$ results | | | Validation results ($p$-values) | | |
|---|---|---|---|---|---|---|---|
| Ensembl ID | Gene Symbol | GWAS SNP(s) | CCmed cis-Gene(s) | CCmed probability | min. GWAS (local eQTLs) | MR-Robin | MultiXcan |
| ENSG00000001461 | NIPAL3 | rs56972983 | WDR55 | 0.888 | $7.0 \times 10^{-4}$ | 0.0421 | 0.3407 |
| ENSG00000007376 | RPUSD1 | rs11693528 | SEPHS1P6 | 0.984 | $9.9 \times 10^{-2}$ | 0.0127 | 0.7957 |
| ENSG00000040487 | PQLC2 | rs7432375 | PCCB | 0.849 | $8.5 \times 10^{-3}$ | 0.0345 | 0.2703 |
| ENSG00000050393 | MCUR1 | rs7523273 | CD46 | 0.974 | $2.7 \times 10^{-2}$ | 0.0316 | 0.0123 |
| ENSG00000064995 | TAF11 | rs8113357 | PRR12 | 0.994 | $3.1 \times 10^{-3}$ | 0.0020 | 0.0073 |
| ENSG00000067177 | PHKA1 | rs8113357 | PRR12 | 0.880 | NA | 0.0069 | NA |
| ENSG00000072756 | TRNT1 | rs56972983 | WDR55 | 0.959 | $4.9 \times 10^{-2}$ | 0.0338 | 0.2067 |
| ENSG00000080345 | RIF1 | rs2905426 | GATAD2A | 0.998 | $1.0 \times 10^{-3}$ | 0.0329 | 0.0761 |
| ENSG00000090054 | SPTLC1 | rs9607771 | SLC25A17 | 0.891 | $3.1 \times 10^{-4}$ | 0.0183 | 0.2751 |
| ENSG00000095906 | NUBP2 | rs7523273 | CD46 | 0.982 | $6.8 \times 10^{-5}$ | 0.0320 | 0.6571 |
| ENSG00000099338 | CATSPERG | rs7085104 | BORCS7 | 0.994 | $1.4 \times 10^{-2}$ | 0.0392 | 0.1363 |
| ENSG00000099810 | MTAP | rs2102949 | PITPNM2 | 0.858 | $7.4 \times 10^{-3}$ | 0.0130 | 0.0624 |
| ENSG00000104886 | PLEKHJ1 | rs679087 | TMTC1 | 0.944 | $1.9 \times 10^{-5}$ | 0.0046 | 0.0011 |
| ENSG00000105583 | WDR83OS | rs301797 | RERE | 0.967 | NA | 0.0193 | NA |
| ENSG00000108559 | NUP88 | rs832187 ; rs832187 | THOC7 ; AC136289.1 | 0.974 ; 0.942 | $3.5 \times 10^{-4}$ | 0.0093 | 0.2180 |
| ENSG00000112667 | DNPH1 | rs679087 | TMTC1 | 0.966 | $3.4 \times 10^{-5}$ | 0.0017 | 0.0028 |
| ENSG00000122490 | PQLC1 | rs832187 | THOC7 | 0.999 | $6.0 \times 10^{-6}$ | 0.0316 | $< 0.0001$ |
| ENSG00000126464 | PRR12 | rs8082590 | DRC3 | 0.973 | $7.1 \times 10^{-7}$ | 0.0011 | $< 0.0001$ |
| ENSG00000127472 | PLA2G5 | rs679087 | TMTC1 | 0.805 | $4.7 \times 10^{-4}$ | 0.0286 | 0.0602 |
| ENSG00000128285 | MCHR1 | rs8113357 | PRR12 | 0.961 | $2.6 \times 10^{-6}$ | 0.0315 | 0.0003 |
| ENSG00000130741 | EIF2S3 | rs7523273 | CD46 | 0.988 | $5.5 \times 10^{-3}$ | 0.0378 | NA |
| ENSG00000130822 | PNCK | rs9607771 | SLC25A17 | 0.845 | $1.4 \times 10^{-2}$ | 0.0077 | NA |
| ENSG00000137142 | IGFBPL1 | rs6434928 | SF3B1 | 0.992 | $9.6 \times 10^{-2}$ | 0.0004 | 0.8936 |
| ENSG00000138778 | CENPE | rs7432375 ; rs7085104 | PCCB ; BORCS7 | 0.900 ; 0.837 | $3.5 \times 10^{-4}$ | 0.0205 | 0.0020 |
| ENSG00000139915 | MDGA2 | rs2905426 | GATAD2A | 0.802 | $5.4 \times 10^{-4}$ | 0.0244 | 0.9456 |
| ENSG00000140497 | SCAMP2 | rs11693528 | SEPHS1P6 | 0.967 | $3.0 \times 10^{-3}$ | 0.0464 | 0.0473 |
| ENSG00000144847 | IGSF11 | rs2905426 | TM6SF2 | 0.867 | $5.0 \times 10^{-2}$ | 0.0480 | 0.0036 |
| ENSG00000145777 | TSLP | rs7523273 | CD46 | 0.998 | $2.4 \times 10^{-3}$ | 0.0324 | 0.0787 |
| ENSG00000146733 | PSPH | rs8082590 | DRC3 | 0.998 | $5.9 \times 10^{-4}$ | 0.0261 | 0.0435 |
| ENSG00000151233 | GXYLT1 | rs7432375 | PCCB | 0.824 | $2.6 \times 10^{-2}$ | 0.0332 | 0.0236 |
| ENSG00000157911 | PEX10 | rs679087 | TMTC1 | 0.937 | $1.9 \times 10^{-2}$ | 0.0252 | 0.0020 |
| ENSG00000162753 | SLC9C2 | rs7085104 | AS3MT | 0.809 | $1.5 \times 10^{-5}$ | 0.0006 | 0.0014 |
| ENSG00000165730 | STOX1 | rs12691307 | INO80E | 0.983 | $1.3 \times 10^{-3}$ | 0.0068 | 0.0079 |
| ENSG00000175264 | CHST1 | rs11693528 | SEPHS1P6 | 0.984 | $4.5 \times 10^{-7}$ | 0.0377 | 0.0142 |
| ENSG00000175826 | CTDNEP1 | rs7523273 | CD46 | 0.999 | $6.5 \times 10^{-5}$ | 0.0321 | 0.0010 |
| ENSG00000177000 | MTHFR | rs56972983 | WDR55 | 0.898 | $1.2 \times 10^{-3}$ | 0.0037 | 0.2608 |
| ENSG00000183628 | DGCR6 | rs11693528 | SEPHS1P6 | 0.984 | $4.2 \times 10^{-2}$ | 0.0067 | 0.7435 |
| ENSG00000184209 | SNRNP35 | rs8082590 | DRC3 | 0.949 | $2.4 \times 10^{-2}$ | 0.0334 | 0.4386 |
| ENSG00000196417 | ZNF765 | rs7523273 | CD46 | 0.810 | $4.4 \times 10^{-2}$ | 0.0445 | 0.5596 |
| ENSG00000196821 | C6orf106 | rs9607771 | SLC25A17 | 0.985 | $1.7 \times 10^{-4}$ | 0.0085 | 0.0044 |
| ENSG00000196937 | FAM3C | rs7432375 | PCCB | 0.960 | $5.4 \times 10^{-3}$ | 0.0077 | 0.0804 |
| ENSG00000196972 | SMIM10L2B | rs9607771 ; rs7523273 | SLC25A17 ; CD46 | 0.944 ; 0.999 | $5.9 \times 10^{-5}$ | 0.0033 | NA |
| ENSG00000197818 | SLC9A8 | rs8082590 | DRC3 | 0.998 | $7.1 \times 10^{-3}$ | 0.0089 | 0.0538 |
| ENSG00000198890 | PRMT6 | rs56972983 | PCDHA4 | 0.961 | $9.0 \times 10^{-3}$ | 0.0215 | 0.1505 |
| ENSG00000204520 | MICA | rs56972983 ; rs11693528 | PCDHA4 ; SEPHS1P6 | 0.933 ; 0.882 | $2.9 \times 10^{-21}$ | 0.0463 | 0.0108 |
| ENSG00000205085 | FAM71F2 | rs12691307 | INO80E | 0.993 | $4.2 \times 10^{-3}$ | 0.0116 | 0.0418 |

#### Description of data used in analyses

##### *The Genotype-Tissue Expression project (GTEx)*

The Genotype-Tissue Expression (GTEx) project is building a comprehensive resource to study tissue-specific gene expression and regulation by collecting postmortem tissue samples from non-diseased tissue sites [4]. Data analyzed in this paper is from GTEx version 8 (v8) [5]. GTEx samples underwent Whole Genome Sequencing at an average coverage of 30X on Illumina HiSeq 2000 or Illumina HiSeq X. GTEx RNA sequencing was performed using the Illumina TrueSeq RNA Sequencing platform. Data was aligned using STAR (v2.5.3a) [6]. Picard [7] was used to mark and remove duplicate reads. Transcripts were quantified using RSEM [8]. RNA-SeQC [9] was used for quality control and gene-level expression quantification, and TMM [10] was used to normalize read counts. Additional details about the genotyping pipeline and sample and variant quality control, and on the RNA-Sequencing pipeline and processing are reported elsewhere [5]. Covariates adjusted for in analyses of GTEx brain tissues included gender, 5 genotype Principal Components, genotyping platform and up to 30 PEER [11] variables.

##### *The Psychiatric Genomics Consortium*

Schizophrenia-risk GWAS statistics were obtained from the second schizophrenia mega-analysis (scz2) conducted by the Psychiatric Genomics Consortium [12]. The GWAS was conducted using up to 36,989 cases and 113,075 controls. In the final analysis, 128 LD-independent SNPs in 108 loci were reported as surpassing the genome-wide significance threshold ($P < 5 \times 10^{-8}$). Additional details of the second PGC GWAS of schizophrenia-risk are reported elsewhere [12].

*The eQTLGen Consortium*

The eQTLGen Consortium performed cis- and trans-eQTL meta-analysis of blood tissue samples from 31,684 individuals across 37 datasets [13]. The cis-eQTL analysis was performed genome-wide while the trans-eQTL analysis was restricted to 10,317 trait-associated variants. After quality control, 16,423 genes were analyzed in the eQTL analyses. Additional details about the eQTLGen Consortium data are reported elsewhere [13].

*The CommonMind Consortium*

The CommonMind Consortium is generating DNA and RNA sequencing, and epigenetic data from ~1000 postmortem brain samples from donors with schizophrenia and bipolar disorder, and from subjects with no neuropsychiatric disorders [14]. RNA sequencing data was generated from dorsolateral prefrontal cortex tissue samples from collections at the Mount Sinai NIH Brain Bank and Tissue Repository, University of Pennsylvania Brain Bank of Psychiatric illnesses and Alzheimer's Disease Core Center, The University of Pittsburgh NIH NeuroBioBank Brain and Tissue Repository, and the NIMH Human Brain Collection Core. Analyses in this paper used Release 1 of the RNA sequencing data from dorsolateral prefrontal cortex samples of people with schizophrenia (N = 258) and control subjects (N = 279). Additional details of CMC data have been reported elsewhere [14].

### Reference

**Author details**
[1]Department of Biostatistics and Informatics, Colorado School of Public Health, University of Colorado Denver, 13001 E. 17th Place, Aurora, Colorado 80045. [2]Department of Public Health Sciences, University of Chicago, 5841 South Maryland Ave MC2000, Chicago, IL 60637. [3] Department of Statistics and Data Science, Carnegie Mellon University, Baker Hall, Carnegie Mellon University, Pittsburgh, PA 15213. [4]Center for Psychiatric Genetics, NorthShore University HealthSystem, 1001 University Place, Evanston, IL 60201. [5] =Department of Psychiatry and Behavioral Neuroscience, 5841 S Maryland Ave, Chicago MC3077, Chicago, IL 60637. [6]Department of Human Genetics, University of Chicago, 920 E 58th St, Chicago, IL 60637.
